# RcsF-independent mechanisms of signaling within the Rcs Phosphorelay

**DOI:** 10.1101/2024.08.29.610257

**Authors:** Anushya Petchiappan, Nadim Majdalani, Erin Wall, Susan Gottesman

**Author notes:** **Corresponding author: Susan Gottesman:**.

## Abstract

The Rcs (regulator of capsule synthesis) phosphorelay is a conserved cell envelope stress response mechanism in enterobacteria. It responds to perturbations at the cell surface and the peptidoglycan layer from a variety of sources, including antimicrobial peptides, beta-lactams, and changes in osmolarity. RcsF, an outer membrane lipoprotein, is the sensor for this pathway and activates the phosphorelay by interacting with an inner membrane protein IgaA. IgaA is essential; it negatively regulates the signaling by interacting with the phosphotransferase RcsD. We previously showed that RcsF-dependent signaling does not require the periplasmic domain of the histidine kinase RcsC and identified a dominant negative mutant of RcsD that can block signaling via increased interactions with IgaA. However, how the inducting signals are sensed and how signal is transduced to activate the transcription of the Rcs regulon remains unclear. In this study, we investigated how the Rcs cascade functions without its only known sensor, RcsF and characterized the underlying regulatory mechanisms for three distinct RcsF-independent inducers. Previous reports showed that Rcs signaling can be induced in the absence of RcsF by a loss of function mutation in the periplasmic oxidoreductase DsbA or by overexpression of the DnaK cochaperone DjlA. We identified an inner membrane protein, DrpB, as a multicopy RcsF-independent Rcs activator in *E. coli*. The loss of the periplasmic oxidoreductase DsbA and the overexpression of the DnaK cochaperone DjlA each trigger the Rcs cascade in the absence of RcsF by weakening IgaA-RcsD interactions in different ways. In contrast, the cell-division associated protein DrpB uniquely requires the RcsC periplasmic domain for signaling; this domain is not needed for RcsF-dependent signaling. This suggests the possibility that RcsC acts as a sensor for some Rcs signals. Overall, the results add new understanding to how this complex phosphorelay can be activated by diverse mechanisms.

**Author summary:** The Rcs phosphorelay signaling cascade regulates the expression of genes related to capsule synthesis, biofilm formation, virulence, and cell division in Enterobacteria and is critical for cell membrane integrity and response to beta-lactam antibiotics and antimicrobial peptides. RcsF is the sole known sensor, but other proteins have been reported to activate this pathway in the absence of RcsF. We have discovered a novel RcsF-independent Rcs activator and found that each of three RcsF-independent proteins activate the system differently. Most significantly, we find that the histidine kinase RcsC can be involved in signal sensing independently of RcsF. Our study sheds light into the complex mechanisms of Rcs activation and adds to our knowledge of non-orthodox signaling systems across organisms.

## Introduction

The Gram-negative bacterial cell wall envelope comprises an outer membrane, the periplasm, a peptidoglycan layer, and an inner membrane [1]. It serves as a protective barrier against environmental insults and a permeability barrier for selective uptake of nutrients. Due to its structural and functional importance for growth and metabolism, bacteria have evolved a multitude of envelope stress response systems to monitor various stresses at the cell wall and respond to them in a timely manner. These systems typically contain a sensor protein which detects the stressor and subsequently activates the downstream signaling pathway leading to changes in gene transcription. A highly conserved envelope stress response pathway in enterobacteria is the Rcs phosphorelay [2]. This cascade responds to outer membrane and peptidoglycan stress from both external and intrinsic sources. While Rcs is a member of the ubiquitous histidine kinase/response regulator signaling systems, in which phosphorylation of the response regulator regulates output, it is significantly more complex than the canonical two-component signaling systems, making it a unique but challenging model to study non-orthodox signaling systems across organisms.

The Rcs pathway has been shown to be activated in response to beta-lactams, antimicrobial peptides, osmotic shock, acid stress, defects in LPS trafficking, peptidoglycan biosynthesis, and contact with a solid surface. The Rcs regulon was first defined for its role in regulating synthesis of capsular polysaccharide but has been shown to include genes related to biofilm formation, motility, virulence, cell morphology, and cell division, among others (reviewed in [2]). How Rcs responds to a wide array of inducers, and the exact mechanism of signal sensing and transduction remains unclear; structural information about most of the components is currently lacking.

The multi-step Rcs phosphorelay comprises an inner membrane hybrid histidine kinase RcsC, an inner membrane phosphotransfer protein RcsD, and the response regulator RcsB (Fig. 1A). The sensor for this cascade is an outer membrane lipoprotein RcsF [3-5]. RcsF detects LPS/peptidoglycan defects and chemical stressors like polymyxin B (presumed to disrupt LPS), A22 (a MreB inhibitor), and mecillinam (a beta-lactam antibiotic) [6, 7]. The phosphorelay is negatively regulated by inner membrane protein IgaA. IgaA is essential and it can only be deleted if the phosphorelay is inactivated by mutations in *rscC*, *rcsD*, or *rcsB* [8]. During normal growth, IgaA represses signaling by interacting with RcsD. When RcsF perceives stress at the envelope, it activates signaling by interacting with the periplasmic domain of IgaA. This relieves the repression of signaling by IgaA, presumably changing the way in which RcsD interacts with RcsC to induce the phosphorelay. A required step in the phosphorelay activation is the autophosphorylation of RcsC. RcsC is a complex histidine kinase, with a receiver domain with a conserved aspartate at its C-terminus (Fig. 1A). A phosphate group is transferred from the conserved histidine residue in RcsC to the aspartate within the RcsC receiver domain. This phosphate group is then transferred to a conserved histidine residue in RcsD and subsequently to an aspartate in RcsB. Phosphorylated RcsB dimers bind promoters to regulate transcription. Alternatively, RcsB can act in concert with other auxiliary partners like RcsA to regulate gene expression (reviewed in [2]). IgaA and RcsD contact each other both in the periplasm and in the cytoplasm; weakening of either interaction leads to activation of the phosphorelay [9]. Thus, IgaA provides a braking mechanism, with the cytoplasmic contact with RcsD as the regulatory switch and the periplasmic interaction preventing Rcs over-activation. The identification of the cytoplasmic interaction as the likely regulatory switch was further underscored by our isolation of a dominant mutation in *rcsD* in its cytoplasmic PAS-like domain (T411A) [9]. The RcsD T411A mutation blocks induction by polymyxin B nonapeptide (PMBN) and other inducers due to tightened cytoplasmic interaction with IgaA.

**Fig. 1:**
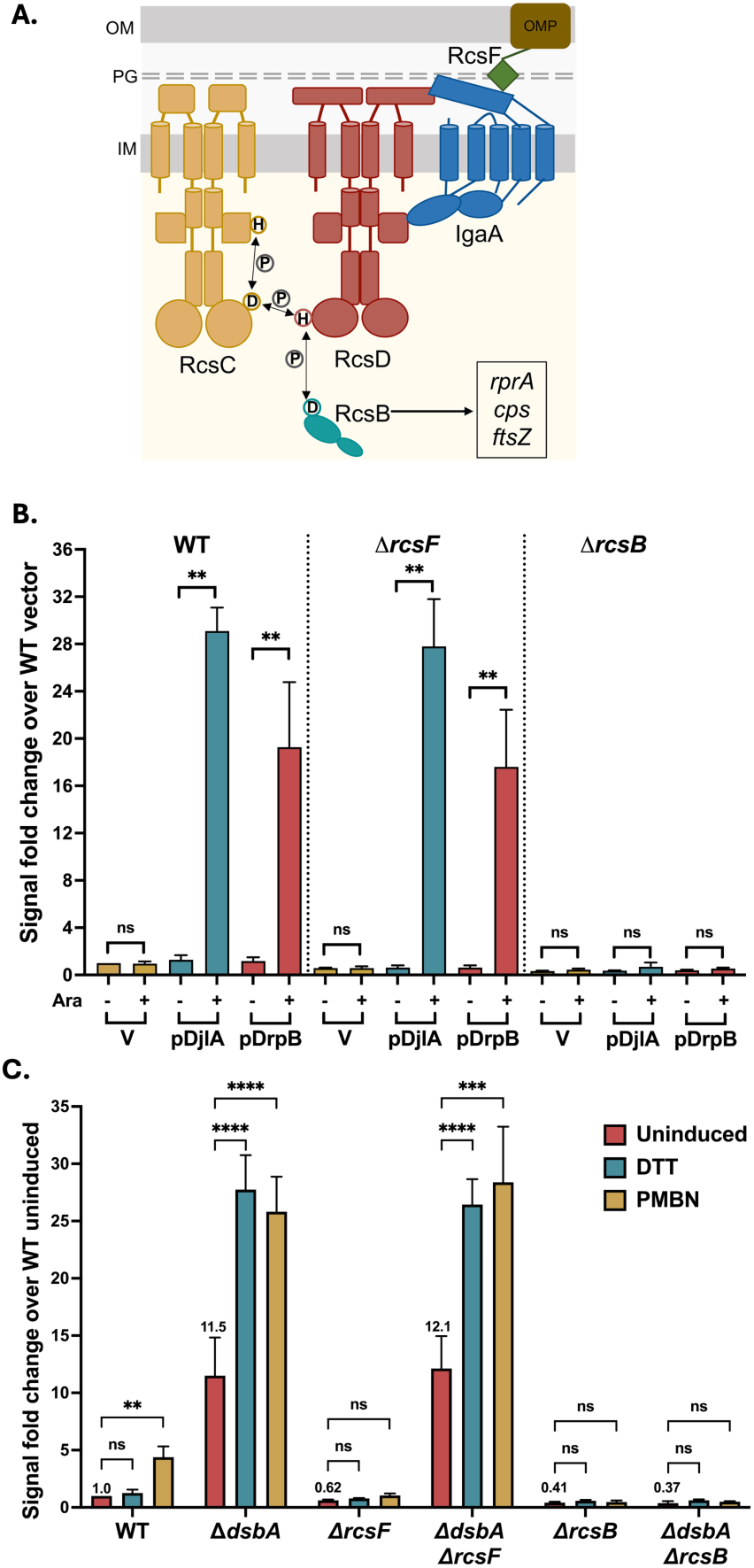
RcsF-independent activators of Rcs signaling. **A. Components of the Rcs phosphorelay:** The Rcs pathway is comprised of the hybrid histidine kinase RcsC, the phosphotransferase RcsD, and the response regulator RcsB as well as the upstream regulatory components RcsF and IgaA. Under non-inducing growth conditions, IgaA represses signaling by interacting with RcsD. RcsF senses damage to the envelope and triggers the RcsC-RcsD-RcsB phosphorelay by interacting with IgaA, altering its interaction with RcsD. **B. Rcs activation by overexpression of DjlA and DrpB:** All strains carry a *rprA* promoter fusion to mCherry (P*_rprA_*::mCherry); mCherry fluorescence acts as an indicator for Rcs activation. For the P*_rprA_*::mCherry assay, the strains overexpressing DjlA (pBAD-DjlA/pPSG961) or DrpB (pBAD-DrpB/pDSW1977) were grown in MOPS minimal glycerol medium containing chloramphenicol (25 μg/ml) and either 0.2% glucose or 0.02% arabinose at 37℃. The RFU at OD 0.4 compared to the uninduced vector control (set to 1) is plotted. The strains used are: WT (EAW8), *rcsF::kan* (AP51), and *rcsB::kan* (EAW31). **C. Rcs signaling in *dsbA* mutants:** For the P*_rprA_*::mCherry assay, the cells were grown in MOPS minimal glucose medium at 37℃. The RFU at OD 0.4 as compared to WT uninduced, set to 1, is depicted here. The cells were treated with either 1mM DTT (blue bars) or 20µg/ml PMBN (brown bars) from the beginning of growth. The strains used were: WT (EAW8), *rcsF::cat* (EAW32), *rcsB::kan* (EAW31), *dsbA::kan* (EAW62), *dsbA::kan rcsF::cat* (EAW67), and *ΔdsbA rcsB::kan* (AP12). Details of the assay are described in Materials and Methods. Data from three independent experiments is plotted as mean with error bars indicating the standard deviation. Values were statistically analyzed using multiple unpaired *t*-tests. Statistical significance is indicated as follows: ns (*P* > 0.05; non-significant), * (*P* < 0.05), ** (*P* ≤ 0.01), *** (*P* ≤ 0.001), and **** (*P* ≤ 0.0001).

While the histidine kinase is the sensor for most two-component systems, RcsC is not known to be a sensor for any of the known Rcs signals. Consistent with this, we previously demonstrated that the RcsC periplasmic domain is dispensable for polymyxin B nonapeptide (PMBN) induction of the Rcs cascade [9]. The outer membrane lipoprotein RcsF remains the only known sensor of the system. RcsF resides within the lumen of the OMP such that a portion of it is surface-exposed [5, 10, 11]. RcsF activates Rcs upon sensing signals perturbing the cell surface and the peptidoglycan layer or upon its mislocalization to the inner membrane.

While the majority of inducing signals require RcsF, there are a number of situations in which RcsF-independent activation of Rcs has been reported. Overexpression of the DnaK cochaperone protein DjlA activates Rcs even in the absence of RcsF [12, 13]. Loss-of-function mutations in *dsbA*, encoding a periplasmic oxidoreductase, also activate the phosphorelay independently of RcsF [4]. Overproduction or mutation of a number of proteins, including TolB [14], DrpB [15], YpdI [16], YmgABC [17], YfgM [18, 19] and YqjA [20], have been linked with Rcs activation but the role of RcsF for signaling has not been reported [4]. Given the complexity of the Rcs phosphorelay, it seems possible that RcsF-independent activation reflects the existence of additional signaling pathways.

Here, we further investigated the nature of RcsF-independent signaling in *dsbA* mutants and upon *djlA* overproduction. We performed a screen for novel RcsF-independent Rcs activators and uncovered a third pathway for RcsF-independent signaling. Comparisons between these pathways delineated three genetically distinct modes of Rcs induction, demonstrating that they are also mechanistically distinct, revealing novel aspects of the regulation of the phosphorelay and implicating the RcsC periplasmic domain in one of these signaling pathways.

## Results

### DrpB is a novel RcsF-independent Rcs activator

With the aim of identifying novel proteins capable of triggering the Rcs cascade in an RcsF-independent manner, we utilized a fluorescence-based genetic screen in *E. coli*. We transformed a plasmid library carrying 3-5 kb fragments of the *E. coli* chromosome into strain EAW34, carrying a reporter for Rcs activation and deleted for *rcsF.* The reporter used is a transcriptional fusion of the *rprA* promoter, an RcsB target, to the mCherry protein, referred to here as P*_rprA_*::mCherry [9, 21]. This strain normally shows low fluorescence in the absence of RcsF, due to low Rcs signaling. The colonies were screened for increased fluorescence and 14 plasmids containing putative RcsF-independent Rcs activators were isolated. Sequencing of the plasmids revealed that 13 of them contained overlapping fragments of the *E. coli* chromosome that included genes encoding the YedR (DrpB) ORF along with the RseX sRNA (Fig. S1A). The remaining plasmid contained genes encoding the YihA protein and the sRNAs Spot 42 and CsrC (Fig. S1B). We previously reported that overexpression of Spot42 induces the Rcs pathway and while that study suggested that much of the Spot 42 effect was RcsF dependent, only a qualitative assay was done [21]. Spot 42, an Hfq-dependent regulatory RNA, regulates translation and stability of multiple mRNAs and has pleiotropic effects in the cell making identification of a single critical target complicated.

To further validate identification of DrpB as capable of RcsF-independent induction, we assayed the growth and fluorescence of wildtype, Δ*rcsF* and Δ*rcsB* strains expressing DrpB or the known RcsF-independent activator DjlA under control of the arabinose-inducible pBAD promoter, using the same P*_rprA_*::mCherry fusion as a reporter for Rcs activation. Overexpression of both DrpB and DjlA showed increased fluorescence, both in the presence and absence of RcsF, confirming their role as multicopy RcsF-independent Rcs activators (Fig. 1B). The increased fluorescence was fully dependent on RcsB for both genes. We did not observe a significant RcsF-independent induction by YihA or *rseX*, and the modest induction by Spot42 in Δ*rcsF* was also not significant (Fig. S1C and S1D). Therefore, we focused our investigation on DrpB. DrpB is a small inner membrane protein and was previously found to activate both the Rcs and the Psp stress response upon overexpression, suggesting that its overexpression leads to membrane stress [15].

We also confirmed that the deletion of *dsbA* activates the Rcs pathway in an RcsF-independent manner, by measuring the fluorescence of the same P*_rprA_*::mCherry reporter in Δ*dsbA,* Δ*dsbA* Δ*rcsF*, and Δ*dsbA* Δ*rcsB* strains (Fig. 1C). Deletion of *dsbA* increased the basal P*_rprA_*::mCherry activity independently of RcsF but not RcsB. Strains devoid of DsbA are defective in disulfide bond formation in periplasmic proteins and DTT treatment is known to exacerbate this problem. Addition of DTT further induced the Rcs pathway, but only in the absence of DsbA. Unexpectedly, we found that PMBN, which normally induces the pathway in an RcsF-dependent-fashion (see WT and Δ*rcsF* results in Fig. 1C), also induced this pathway, to a similar extent as DTT, in both 11*dsbA* and 11*dsbA* 11*rcsF* strains (Fig. 1C). Together, these results provide evidence for three RcsF-independent Rcs signaling situations – lack of DsbA and overproduction of DjlA or DrpB.

We next investigated whether these inducers of Rcs signaling depend on each other. We first measured the Rcs signaling in *dsbA* mutants in cells also deleted for the chromosomal copies of *djlA* or *drpB* (Fig. S2A). Signaling in the 11*dsbA* 11*djlA* and 11*dsbA* 11*drpB* strains was essentially unchanged from that seen in the Δ*dsbA* strain (see Fig. 1C). Therefore, the absence of DsbA does not signal by mimicking overexpression of DjlA or DrpB. In addition, overexpression of DjlA or DrpB induces signaling in a *dsbA* deletion strain (Fig. S2B), although the overexpression of DrpB led to poor growth in the absence of DsbA (see OD values, Fig S2B). We also tested if the multicopy activators DjlA and DrpB require each other to trigger Rcs signaling and found that they did not (Fig. S2C). Together, these results indicate that DsbA, DjlA, and DrpB do not require one another for signaling and possibly employ different mechanisms of Rcs activation. We next investigated the pathways they each use for RcsF-independent activation of Rcs.

### DrpB has a unique requirement for the RcsC periplasmic domain for signaling

The Rcs pathway can be activated simply by increasing the levels of RcsB [22]. If this were how the RcsF-independent factors were working, they might be independent of the histidine kinase, RcsC and the phosphotransfer protein RcsD. Previous work had demonstrated that multicopy DjlA and DrpB are still dependent on RcsC for signaling [12, 13, 15]. As seen in Fig. 2, while deletion of *rcsC* caused an increase in the basal level of P*_rprA_*::mCherry expression, as previously seen [9, 21], there was no further induction in the absence of RcsC for any of these factors.

**Fig. 2:**
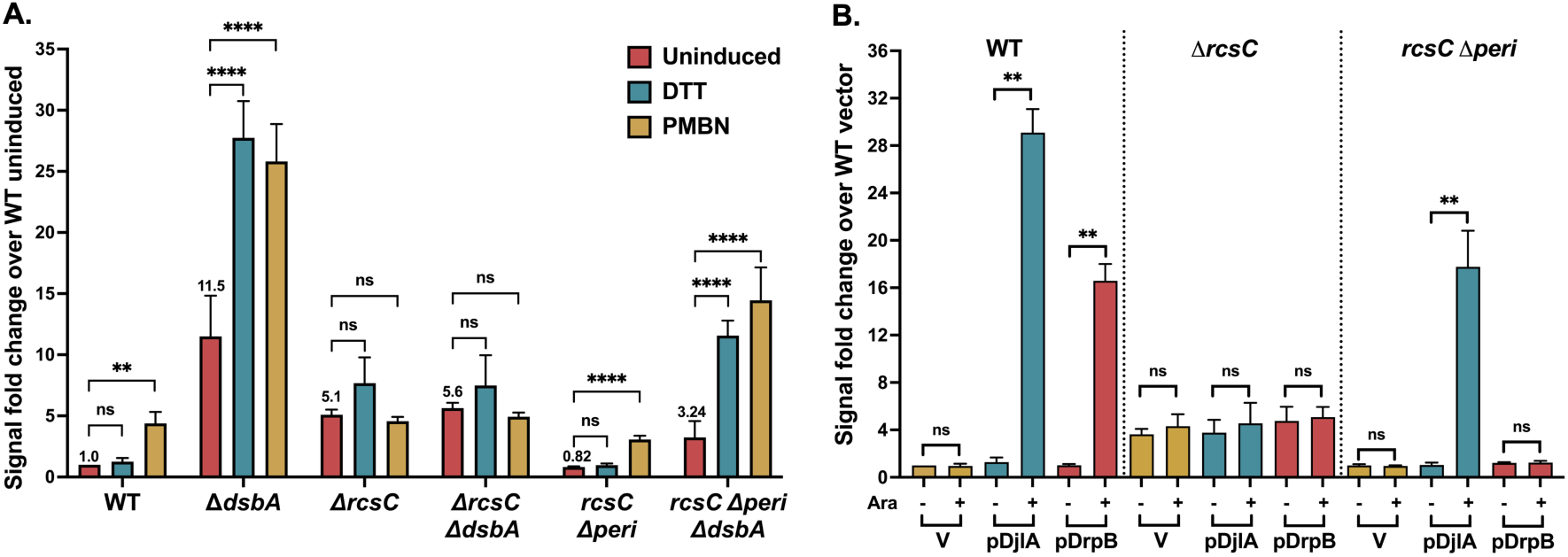
Requirement of RcsC periplasmic domain for RcsF-independent signaling by DrpB. **A. *dsbA* mutants do not need the RcsC periplasmic domain for signaling:** For the P*_rprA_*::mCherry assay, the cells were grown in MOPS minimal glucose medium at 37℃. The RFU at OD 0.4 as compared to WT uninduced, set to 1, is depicted here. The cells were treated with either 1mM DTT or 20µg/ml PMBN. **B. DrpB, but not DjlA, requires the RcsC periplasmic domain for Rcs activation:** For the P*_rprA_*::mCherry assay, the strains overexpressing DjlA (pBAD-DjlA/pPSG961) or DrpB (pBAD-DrpB/pDSW1977) were grown in MOPS minimal glycerol medium containing chloramphenicol (25 μg/ml) and either 0.2% glucose (-Ara) or 0.02% arabinose at 37℃. The RFU at OD 0.4 compared to the uninduced vector control is plotted. The strains used were: WT (EAW8), *dsbA::kan* (EAW62), *rcsC::tet* (EAW18),), *rcsC::tet dsbA::kan* (EAW63), *rcsC Δperi* (EAW70), and *dsbA::kan rcsC Δperi* (EAW74). Values (mean ± SD) were statistically analyzed using multiple unpaired *t*-tests. Statistical significance is shown as: ns (*P* > 0.05; non-significant), ** (*P* ≤ 0.01), and **** (*P* ≤ 0.0001).

We previously showed that the RcsC periplasmic domain is not needed for the RcsF-dependent PMBN response [9]. We assayed the Rcs activation upon deletion of *dsbA* or overexpression of DjlA or DrpB in strains encoding a chromosomal version of RcsC missing its periplasmic domain (*rcsC* 11*peri*). Fig. 2A shows that PMBN was able to induce signaling in the *rcsC* 11*peri* strain as previously found. Deletion of *dsbA* in the *rcsC* 11*peri* strain increased the basal level of signaling, although not to the extent seen for the *rcsC*^+^ strain (Δ*dsbA*); DTT or PMBN further increased signaling (Fig. 2A). These results demonstrate that the effect of Δ*dsbA* is independent of the RcsC periplasmic region. DjlA overexpression was also able to activate signaling in the *rcsC* 11*peri* strain (Fig. 2B). However, upon DrpB overexpression, no Rcs activation was observed in the *rcsC* 11*peri* strain (Fig. 2B). This is the first evidence of a role for the RcsC periplasmic domain in signaling and provides clear evidence that DrpB acts in a manner that is distinct from DjlA overexpression or loss of DsbA.

### DjlA, but not DsbA and DrpB, can induce Rcs signaling in a RcsD T411A mutant

As for deletion of *rcsC* (Fig. 2), all three factors were unable to induce signaling in the absence of the phosphorelay protein RcsD (Fig. 3). RcsD T411A is an allele of RcsD expressing a mutant in the cytoplasmic domain of RcsD that has a higher affinity for IgaA, blocking RcsF-dependent PMBN signaling [9] (Fig. 3A). Strains carrying *rcsD* T411A were used to further explore the role of IgaA-RcsD interactions in RcsF-independent signaling. DsbA signaling was blocked by the *rcsD* T411A mutation; the basal level was reduced to that seen in *rcsD* T411A alone, and the DTT or PMBN increase was blocked (Fig. 3A). Overexpression of DrpB also could not induce signaling in the *rcsD* T411A strain (Fig. 3B). However, DjlA overexpression activated signaling in this strain, indicating its unique ability to overcome the effect of the *rcsD* T411A mutation.

**Fig. 3:**
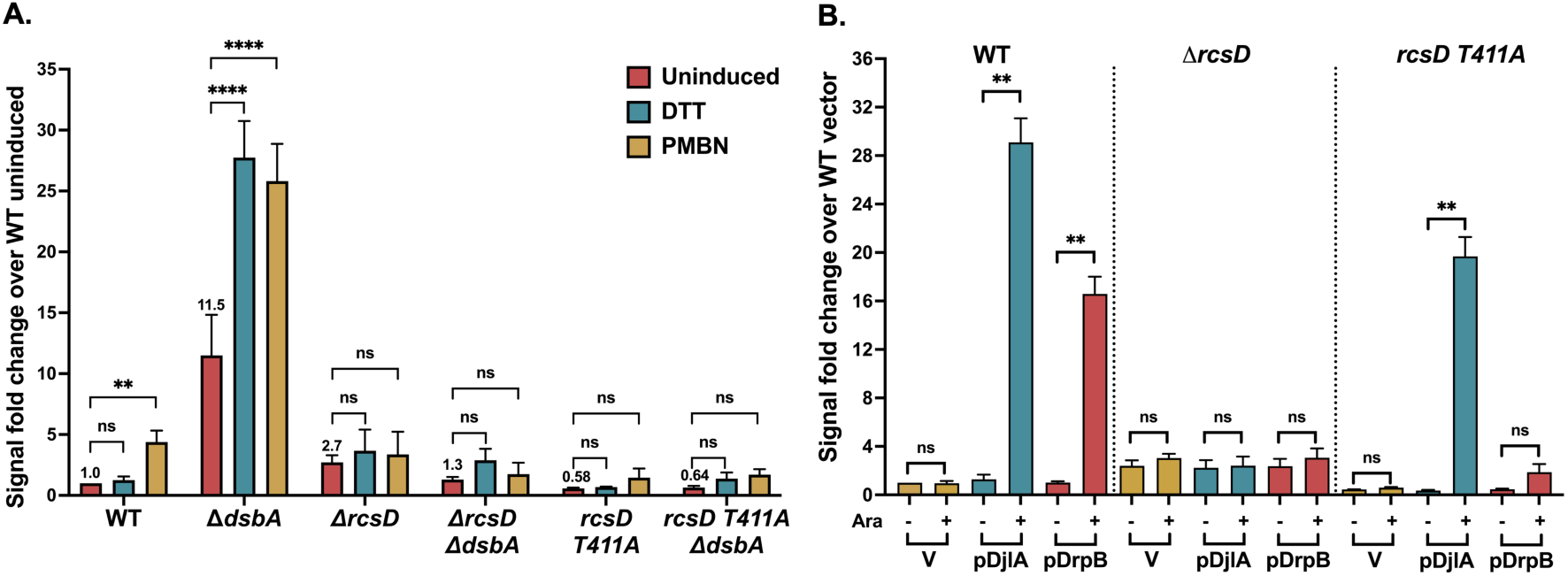
Effect of RcsD T411A mutation on RcsF-independent signaling. **A. RcsD T411A mutation blocks signaling by *dsbA*:** For the P*_rprA_*::mCherry assay, the cells were grown in MOPS minimal glucose medium at 37℃. The RFU at OD 0.4 as compared to the uninduced WT control, set to 1, is depicted here. The cells were treated with either 1mM DTT or 20µg/ml PMBN. Details of the assay are described in Materials and Methods**. B. DjlA, but not DrpB, can overcome the RcsD T411A mutation for Rcs activation:** For the P*_rprA_*::mCherry assay, the strains overexpressing DjlA (pDjlA/pPSG961) or DrpB (pDrpB/pDSW1977) were grown in MOPS minimal glycerol medium containing chloramphenicol (25 μg/ml) and either 0.2% glucose or 0.02% arabinose at 37℃. The RFU at OD 0.4 compared to the vector uninduced control, set to 1, is plotted. The strains used were: WT (EAW8), *dsbA::kan* (EAW62), *ΔrcsD* (EAW19), *ΔdsbA rcsD541::kan* (AP13), *rcsD T411A* (EAW121), and *rcsD T411A dsbA::kan* (AP14). Statistical significance was calculated using multiple unpaired *t*-tests and is shown as follows: ns (*P* > 0.05; non-significant), * (*P* < 0.05), ** (*P* ≤ 0.01), and **** (*P* ≤ 0.0001).

We considered the possibility that DjlA could function in the *rcsD* T411A strain because it was the strongest inducer of the conditions tested. As seen in Fig. 1B and 1C, DjlA overexpression gave around 30-fold induction, which is higher than DrpB overexpression (∼17 fold), DsbA deletion (∼13 fold), and the RcsF-dependent response to PMBN (∼4 fold). As a test of this model, we examined the ability of RcsD T411A to block signaling when RcsF is highly active. Overproduction of RcsF or expression of a derivative (RcsF^IM^, carrying mutations S17D and M18Q) that localizes the protein to the inner member and constitutively activates Rcs signaling [7, 23] both induce Rcs signaling, with the inner-membrane localized mutant RcsF derivative, RcsF^IM^, being the stronger activator, inducing better than 25-fold (Fig.S3). However, the T411A mutation was still able to block this signaling (Fig. S3). These results suggest that DjlA is unique in its ability to fully overcome the *rcsD* T411A allele.

Thus, each of the three RcsF-independent factors has distinct patterns of induction, suggesting that they act in different manners. While all three are dependent upon RcsC and RcsD, only DrpB signaling depends on the RcsC periplasmic domain and only DjlA can bypass the *rcsD* T411A allele to induce signaling. Below we further examine how each of these factors works.

### DsbA is needed for efficient IgaA-RcsD periplasmic interactions

DsbA resides in the periplasm and plays a central role in disulfide bond (DSB) formation in *E. coli* and other bacteria [reviewed in [24]]. DsbA, in concert with DsbB, introduces disulfide bonds into newly synthesized periplasmic proteins, predominantly between consecutive cysteine residues [25-27]. *dsbA* null mutants show a pleotropic phenotype consistent with defects in the cell envelope - they are mucoid, non-motile, sensitive to DTT as well as some drugs, show decreased fitness and attenuated virulence [24, 25, 28, 29]. Periplasmic and membrane proteins that require disulfide bonds frequently require DsbA for proper folding. One such protein is the Rcs sensor protein, the outer membrane lipoprotein RcsF [30]. RcsF has two non-consecutive disulfide bridges and impaired disulfide bond formation leads to its degradation [31, 32]. Thus, RcsF is non-functional in a *dsbA* mutant, supporting the idea that mucoidy in the *dsbA* mutant strain, likely reflecting Rcs induction, cannot be due to activation via RcsF; this is confirmed by the induction of the P*_rprA_*::mCherry reporter in the absence of RcsF in Fig. 1C. This is further validated by the absence of Rcs induction in response to Mecillinam or the MreB inhibitor A22 by the *dsbA* mutants, both dependent on RcsF (Fig. S4A). Note that this result also distinguishes PMBN from these other inducers as uniquely still able to induce the Rcs phosphorelay in the *dsbA* mutant (Fig. 2A, Fig. S4B; compare differences in fold induction).

Aside from RcsF, RcsC, RcsD, and IgaA each have a periplasmic domain that could possibly be targets for DsbA. RcsD does not have any conserved cysteine residues within its periplasmic domain, ruling it out as a direct target of DsbA.

RcsC has two conserved cysteine residues in its periplasmic domain, but since the *11dsbA rcsC* 11*peri* strain is still DTT-inducible (Fig. 2A), this cannot be the critical target for DsbA. However, our results do suggest that, directly or indirectly, the RcsC periplasmic domain contributes to phosphorelay activation in the absence of DsbA. While deletion of the RcsC periplasmic domain had only a modest effect on P*_rprA_*::mCherry basal level or induction in a *dsbA*^+^ strain (compare WT to *rcsC*11*peri* data, Fig. 2A), levels of expression in the Δ*dsbA* mutants were significantly lower for 11*dsbA rcsC* 11*peri* than for 11*dsbA rcsC*^+^ (Fig. 2A), with Δ*dsbA* 12-fold higher than WT in the absence of induction, while 11*dsbA rcsC*11*peri* was only 3.2-fold increased compared to WT or *rcsC*11*peri* (Fig. 2A). To determine if this reduction in induction was dependent upon RcsC disulfide bonds, we examined the behavior of a strain expressing RcsC mutant for the C111 and C154 periplasmic cysteine residues (*rcsC C111A C154A*, named here as *rcsC C2A*) in *dsbA*^+^ or *ΔdsbA* strains. As seen in Fig. S4B, the *rcsC C2A dsbA*^+^ strain has the same pattern of expression as the WT strain or a *rcsC* 11*peri* strain (Fig.2A), responding to PMBN but not DTT. The *11dsbA rcsC C2A* strain, unlike the *rcsC*11*peri* strain, responds to both DTT and PMBN and this response is not muted as in *rcsC*11*peri* (compare to Fig. 2A). This suggests the effect of the *rcsC* 11*peri* does not reflect a role of the RcsC periplasmic cysteine residues and is likely more indirect; this was not further explored here.

IgaA has four conserved periplasmic cysteine residues, suggesting that there are two disulfide linkages in its periplasmic domain. A study in *Salmonella* supported the idea that at least the first of the disulfide bonds was important for proper IgaA function [33]. Given that IgaA is essential, the *dsbA* mutant cannot lead to fully non-functional IgaA. However, if the disulfide bonds are not properly formed, the periplasmic interaction between IgaA and RcsD could be affected in the absence of DsbA, leading to the observed Rcs activation. We have previously shown that IgaA and RcsD interact in a bacterial two hybrid (BACTH) assay, and that the interaction detected by this assay is primarily dependent upon a periplasmic contact between the proteins [9]. This assay is based on the reconstitution of the Bordetella adenylate cyclase from its two fragments, T18 and T25, in an adenylate cyclase mutant (*cya*) strain BTH101 [34, 35]. Fusion of each of these fragments to an interacting protein reconstitutes adenylate cyclase, assayed by the Cya-dependent synthesis of beta-galactosidase. Both orientations, IgaA-T18/RcsD-T25 as well as IgaA-T25/RcsD-T18, have been demonstrated to show a strong interaction ([9] and Fig. 4A, WT strain).

**Fig. 4:**
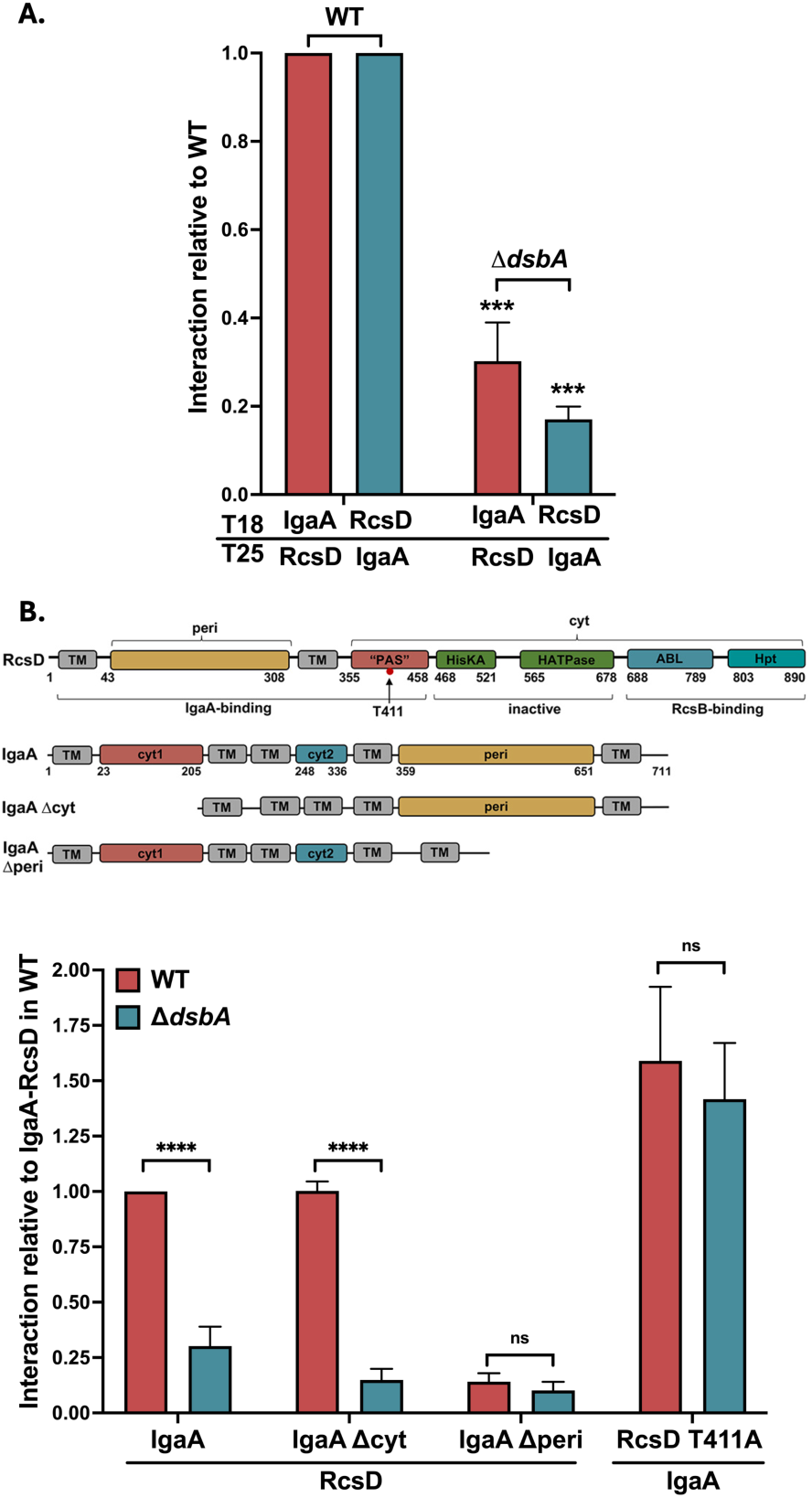
DsbA is needed for efficient IgaA-RcsD periplasmic interactions. **A. IgaA-RcsD interactions are weakened in Δ*dsbA*.** Beta-galactosidase activity was measured in a Δ*cyaA* strain (BTH101 or AP 58 (BTH101 Δ*dsbA*)) expressing two plasmids encoding the T18 and T25 domains of adenylate cyclase fused to the proteins of interest and the expression measured compared to vector background. The IgaA/RcsD protein fusion plasmids paired with their cognate vector had very low activity; the three controls were averaged and used as “background” for normalization. The IgaA-RcsD interaction was normalized to 1 and other interactions were plotted relative to this interaction in WT. This data is compiled from separate sets of assays, each normalized relative to the IgaA/RcsD signal in that experiment. The P values are relative to the *dsbA*^+^ values. **B. Periplasmic interactions are affected in Δ*dsbA* and a T411A mutation strengthens this IgaA interaction.** IgaA and RcsD/T411A were fused to the T18 and T25 domains, respectively. The IgaA-RcsD interaction in WT was normalized to 1 and all other interactions are plotted relative to this interaction. The RcsD T411A mutation tightens the IgaA-RcsD interaction in WT as well as Δ*dsbA*. The plasmids used were pEAW1 (IgaA-T18), pEAW8 (RcsD-T25), pEAW2 (IgaA-T25), pEAW7 (RcsD-T18), pAP101 (IgaA Δcyt), pEAW1peri (IgaA Δperi), and pEAW8T (RcsD T411A-T25). Values (mean ± SD) from independent experiments were statistically analyzed using multiple unpaired *t*-tests. Statistical significance is shown as follows:ns (*P* > 0.05; non-significant), *** (*P* ≤ 0.001), and **** (*P* ≤ 0.0001).

We introduced the *dsbA* null mutant into the host for the BACTH assay and studied its effect on the IgaA-RcsD interaction, compared to the WT strain (Fig. 4A). The interaction between IgaA and RcsD was significantly impaired in the *dsbA* mutant, suggesting that DsbA is needed to maintain a proper IgaA-RcsD interaction, possibly due to its role in IgaA folding. Given the localization of DsbA in the periplasm, we would expect that it is the periplasmic interaction of IgaA with RcsD that should be defective in the absence of DsbA. This was confirmed first by testing the interaction of RcsD with various domain deletion constructs of IgaA. IgaA has five transmembrane helices anchoring two cytoplasmic domains and a large periplasmic domain (schematic in Fig. 4B). The IgaA 11*peri* (11384-649) protein has been shown to have a weak interaction with RcsD, confirmed here; loss of DsbA did not further decrease the interaction (Fig. 4B). Deletion of the two cytoplasmic domains (IgaA 11cyt), on the other hand, did not decrease the interaction, but was very sensitive to loss of DsbA (Fig. 4B). Neither of these deletions significantly affected the levels of expression of the IgaA-T18 protein, in the presence of absence of DsbA (Fig. S4D). This data is best explained by a change in the IgaA periplasmic domain in the absence of DsbA, reducing the periplasmic interaction with RcsD.

A crystal structure of the *E. coli* IgaA periplasmic domain shows the presence of the two disulfide linkages to be C404-C425 and C498-C504, consistent with previous work [33, 36]. We mutated the Cysteine residues to Serine and tested their effect on the IgaA-RcsD interaction, either in pairs (C404S C425S or C498S C504S) or in a mutant with all four mutated (C4S) (Fig. S4C). Breaking either linkage significantly weakened the interaction of IgaA with RcsD. As with the data for Δ*dsbA* in Fig. 4B, the RcsD T411A allele overcame much (but not all) of the loss of interaction in the IgaAC4S derivative (Fig. S4C). Therefore, for this interaction assay, C4S mimics the effect of loss of *dsbA*, supporting the idea that the disulfide linkages are required for proper folding of the periplasmic domain of IgaA and its periplasmic interaction with RcsD. Mutations in IgaA may lead to its degradation during the stationary phase [33] but the overexpressed IgaA C4S mutant was stable in the T18 fusion, suggesting DsbA primarily affects its folding (Fig. S4D).

The prediction from these interaction assays is that mutations of the IgaA cys residues should also lead to increased activation of the Rcs phosphorelay, as seen for Δ*dsbA* in the presence of DTT. The chromosomal copy of *igaA* was replaced with *igaA*C4S in an *rcsD* mutant strain carrying the P*_rprA_*::mCherry reporter. While this strain (AP168) is not fluorescent and is non-mucoid, due to lack of RcsD, when a plasmid carrying RcsD was introduced into the cells, colonies became mucoid, consistent with activation of the Rcs phosphorelay. Growth was poor and it seems likely that this C4S strain, like other strains that express the Rcs phosphorelay at high levels, rapidly picks up suppressing mutations. Growing the transformed colonies without further purification in LB and assaying the P*_rprA_*::mCherry signal demonstrated that the signal was higher than that seen with the Δ*dsbA* strain and was not affected by DTT (Fig. S4E). Thus, blocking disulfide bond formation in IgaA via mutation of the cysteine residues induces Rcs signaling and mimics the Δ*dsbA* phenotype. Because either pair of cysteine mutants abrogates the full interaction with RcsD (Fig. S4C), it seems likely both disulfide bonds are important for proper IgaA folding.

### DjlA can act as a cochaperone and mediate IgaA-RcsD cytoplasmic interactions

DjlA is the third member of the DnaJ domain family in *E. coli*. Proteins with this domain, including DnaJ and CbpA in *E. coli*, collaborate as part of the widely conserved DnaK/Hsp70 chaperone network. DjlA is localized at the cytoplasmic membrane with a short N-terminal region in the periplasm followed by a single transmembrane helix [37]. The conserved J-domain, needed for DnaK interaction, is in the cytoplasm at the C-terminal. A TerB-like (tellurite resistance) domain of unknown function lies between the J-domain and the membrane helix. The role played by this domain has not been clearly defined but it has been speculated to be involved in membrane/lipid-interaction, stress-sensing, DNA processing, and phage defense [38, 39]. Although the precise cellular function and substrates of DjlA are unknown, it is a bonafide DnaK cochaperone capable of assisting DnaK in protein refolding [40]. DjlA is not essential for growth and a null mutant did not reveal any significant phenotype except a delayed onset of mucoidy at low temperature [13]. On the other hand, overproduction of DjlA was clearly toxic, with the cells exhibiting growth defects suggestive of a role of DjlA in cell division or membrane integrity [37]. Furthermore, strains overexpressing DjlA demonstrated increased capsule synthesis, as result of Rcs activation, and hypersensitivity to certain drugs and chelators [41]. The J-domain is not strictly required for the increased drug sensitivity; however, it is essential for its role as a cochaperone and Rcs activator [13]. DjlA-dependent activation of Rcs signaling requires DnaK and the nucleotide release factor GrpE, but not DnaJ or CbpA [13]. A functional J-domain and the transmembrane helix were both reported to be essential for Rcs induction by DjlA [13, 42, 43].

As a membrane-anchored cochaperone, it is possible that DjlA directs DnaK to modulate the folding and/or conformation of an Rcs component or the interaction of Rcs components. This is further supported by our observation that DjlA, uniquely, can overcome the tight interaction between IgaA and RcsD T411A (Fig. 3B). Therefore, we tested the effect of DjlA overproduction on the IgaA-RcsD interaction in BACTH assays. We examined the IgaA-RcsD interaction when *djlA* was cloned downstream of IgaA-T18, such that its expression is controlled by the same IPTG-inducible promoter as IgaA. Fig. 5A shows that there is a marked decrease in the IgaA-RcsD interactions when DjlA is overexpressed. To confirm that this effect is due to the cochaperone activity of DjlA, we also tested an inactive DjlA mutant, a H233Q mutation known to be defective for DnaK cochaperone activity [13, 40]. The overexpression of this DjlA H233Q mutant did not disrupt the IgaA-RcsD interaction. Next, we asked if DjlA can weaken the tight interaction between IgaA and the RcsD T411A mutant (Fig. 5B). DjlA overcame this strong interaction and could act even when the periplasmic domain of RcsD was not present (Fig. 5B, T411A 11peri). This contrasts with loss of DsbA, which affected the periplasmic but not the cytoplasmic interactions between IgaA and RcsD (Fig. 4), and is consistent with the ability of overproduced DjlA to cause Rcs induction even when RcsDT411A is present (Fig. 3B). We also tested if any of the other Rcs components are required for DjlA to affect these interactions by carrying out the same assay in derivatives of the BTH101 strain carrying 11*rcsB* (EAW1), Δ*rcsC* (EAW2)*, or* Δ*rcsF* (EAW4) mutants. DjlA was able to weaken the interaction between IgaA and RcsD in the absence of each of these proteins (Fig. 5C). This suggests that DjlA acts directly as a cochaperone, remodeling the RcsD-IgaA interaction with one or both proteins as direct targets for DjlA.

**Fig. 5:**
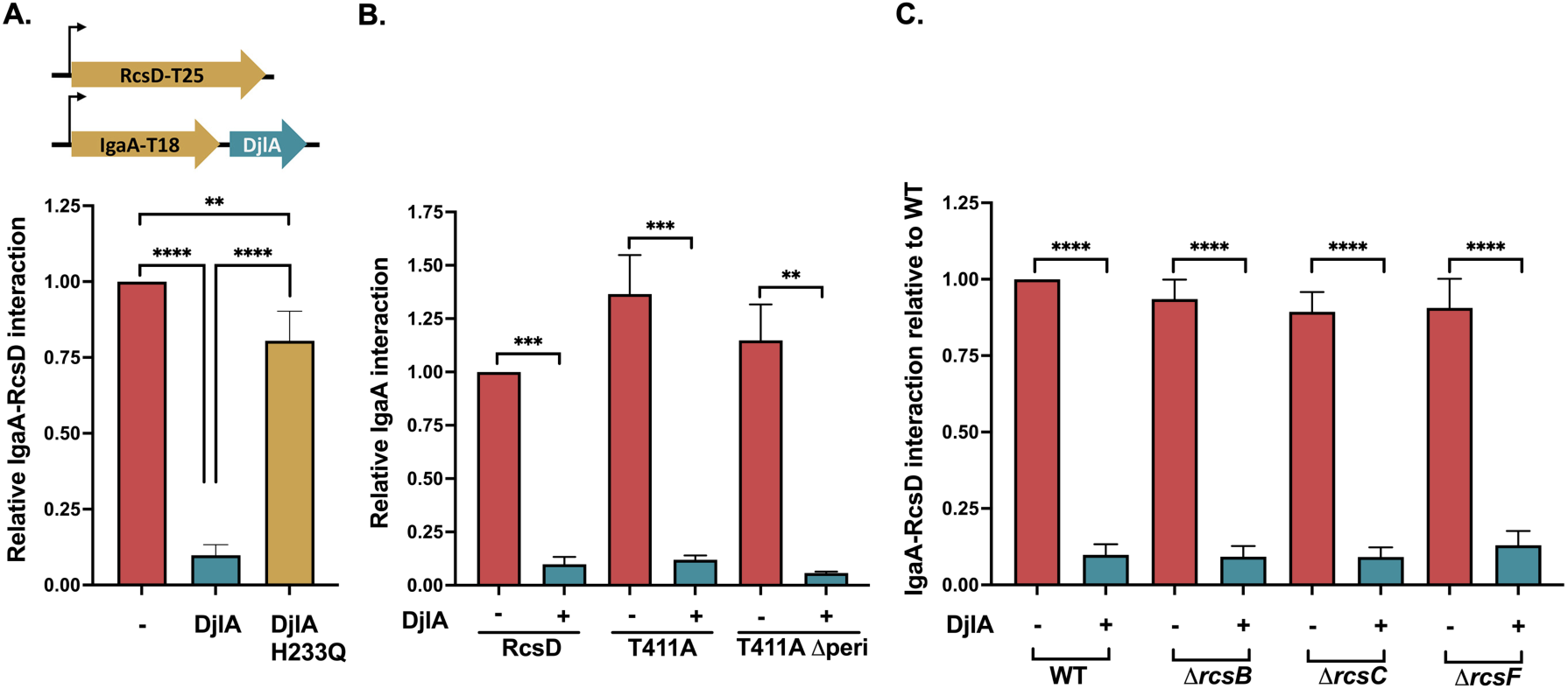
DjlA weakens IgaA-RcsD interactions. **A. DjlA loosens IgaA-RcsD interactions:** Beta-galactosidase activity was measured in a Δ*cyaA* strain (BTH101) in the presence of the indicated plasmids. IgaA and RcsD/RcsD T411A were fused to the T18 and T25 domains, respectively. DjlA or the H233Q mutant of DjlA, inactive as a co-chaperone, was cloned downstream of IgaA under the same promoter control. The IgaA-RcsD interaction was normalized to 1 and all other interactions are plotted relative to this interaction. **B. DjlA can disrupt cytoplasmic interactions of IgaA and RcsD:** The IgaA-RcsD interaction was normalized to 1 and all other interactions are plotted relative to this interaction**. C. RcsB, RcsC, RcsF are not needed for DjlA to weaken interactions.** The IgaA-RcsD interaction was tested in BTH101 (WT), EAW 1 (*BTH 101 rcsB::tet*), EAW2 (*BTH 101 rcsC::tet*) and EAW4 (*BTH 101 rcsF::cat*). Plasmids used in this set of experiments were pEAW1 (IgaA-T18), pEAW8 (RcsD-T25), pEAW8T (RcsD T411A-T25), pAP804 (RcsD T411A Δperi -T25), pAP1401 (IgaA-T18 + DjlA), and pAP1402 (IgaA-T18 + DjlA H233Q). Statistical significance was calculated using multiple unpaired *t*-tests and is shown as follows: ** (*P* ≤ 0.01), *** (*P* ≤ 0.001), and **** (*P* ≤ 0.0001).

Previous studies speculated that RcsC is the target of DjlA likely via transmembrane domain interactions, although that work was done before RcsD and IgaA were known or characterized [42]. As shown above, the periplasmic domain of RcsC was not needed for DjlA induction, but the role of the RcsC transmembrane helices had not been tested. We had previously demonstrated signaling by a RcsC_MalF_ strain containing the RcsC cytoplasmic domain fused to the first two transmembrane (TM) helices of the maltose-transporter membrane protein MalF [9]. RcsC_MalF_ increased the basal Rcs signal of the strain, but it responded to a RcsF-dependent signal like PMBN [9]. DjlA overexpression activated Rcs signaling in this strain, demonstrating that DjlA does not need to interact with the RcsC TM helices (Fig. S5A).

We further examined the role of the DjlA TM helix in DjlA-dependent Rcs activation. Point mutants in the TM helix of DjlA have been shown to completely block Rcs activation without affecting its stability or localization [42]. Furthermore, the DjlA TM helix is a dimerization domain and contains conserved glycine residues important for Rcs induction [44]. We replaced the DjlA TM helix with the first two MalF TM helices such that the cytoplasmic portion of DjlA is anchored to the membrane with the MalF helices (DjlA_MalF_; Fig. S5B). We tested P*_rprA_*::mCherry induction upon overexpression of this DjlA_MalF_ variant in a 11*djlA* 11*rcsF* strain. We observed that this chimeric DjlA_MalF_ variant is capable of inducing Rcs signaling, unlike the DjlA H233Q mutant or the DjlA ΔTM variant lacking any TM helix (Fig. S5B). These results lead us to conclude that the DjlA TM helix does not have an active role in Rcs induction, but that membrane localization is indeed essential. Previous reports did not observe any Rcs induction with either DjlA 11TM or a MalF-DjlA chimeric variant [13, 42, 44]. The discrepancy could be due to the somewhat different DjlA_MalF_ protein construct used by us or may reflect the increased sensitivity of our P*_rprA_*::mCherry fluorescent reporter as compared to the *cps-lacZ* beta-galactosidase assay used previously. We further tested the ability of DjlA 11TM and DjlA_MalF_ to break the IgaA-RcsD interaction using the BACTH assay. Fig. S5C shows that both of these DjlA constructs can disrupt the interaction. This is consistent with the DjlA TM helix not having a specific role in recognizing its Rcs target protein. T18-IgaA levels did not decrease upon overexpression of DjlA (Fig. S5D). Surprisingly, DjlA 11TM was also capable of disrupting the IgaA-RcsD interactions in BACTH even though it was not functional in Rcs activation. Possibly this is due to the difference in protein levels of IgaA/RcsD/DjlA between these two assays. Thus, the TM helix of DjlA is important for inducing signaling but may be less important for recognizing its Rcs substrate.

DjlA was also able to disrupt the interaction of IgaA11cyt with RcsD (Fig. S5C). We have previously shown that the periplasmic interaction of IgaA and RcsD drives the signal in the BACTH assay [9]. Assuming DjlA, with the J-domain in the cytoplasm, interacts with cytoplasmic domains of its substrates, the observation that full-length DjlA can disrupt IgaA11cyt-RcsD interactions may suggest that DjlA affects interactions of IgaA with the PAS-like domain of RcsD, leading to conformational changes that lessen the strong periplasmic contacts between RcsD and IgaA. Interestingly, the DjlA 11TM was less effective than full length DjlA in disrupting IgaA11cyt-RcsD interactions.

If multicopy DjlA weakens IgaA-RcsD interactions, it could be speculated that it has a role in modulating these interactions even in single copy under appropriate physiological conditions. However, we failed to find a single copy phenotype of Δ*djlA* for Rcs signaling. A strain devoid of DjlA responded like a wild-type strain to varying doses of PMBN (Fig. S6A) and responded to A22 and mecillinam (Fig. S6B). It also responded similarly even at elevated temperature (42°C) (Fig. S6C). While we were unable to detect a role for single copy DjlA for Rcs induction, BACTH assays of IgaA-RcsD interactions showed a somewhat better interaction in the absence of DjlA, consistent with its proposed role in chaperoning this interaction (Fig. S6D). Consistent with a role for DjlA for the cytoplasmic interaction of these proteins, an increased interaction of RcsD Δperi with IgaA was also seen in the Δ*djlA* strain. RcsD T411A, with a tighter cytoplasmic interaction with IgaA, was unaffected by the absence of DjlA. These results are consistent with the idea that DjlA plays a role in the cytoplasmic signaling interactions of IgaA and RcsD, but under conditions not yet identified.

### DrpB activates Rcs independently of its cell division role

In contrast with DsbA and DjlA, induction by DrpB overexpression has a requirement for the RcsC periplasmic domain for Rcs activation (Fig. 2). Whether there is a direct interaction between DrpB and RcsC for the signaling to occur or the RcsC periplasmic domain senses a signal generated by DrpB overexpression is not yet known. DrpB is a small (100 aa) protein with two transmembrane helices and a small periplasmic region (see schematic in Fig. S7A). There are no additional recognizable domains. However, DrpB was identified as a multicopy suppressor of the growth defect of a Δ*ftsEX* mutant at low ionic strength [45]. FtsEX is an ABC transporter that coordinates peptidoglycan remodeling and cell division, allowing cell separation. FtsEX helps in recruitment of downstream cell division proteins to the Z-ring, cell constriction, and activation of periplasmic peptidoglycan amidases[46-48]. FtsEX mutants display chaining defects and have low viability in low-osmolarity medium. DrpB localizes to the divisome and overexpression allowed the Δ*ftsEX* [45]. Activation of amidases by FtsEX is not strictly required for survival in low salt medium so it is likely that overproduction of DrpB rescued the growth defect by improving recruitment of downstream components of the divisome [45, 48]. Although no cell division or cell morphology defects were observed in a Δ*drpB* strain, a double deletion strain with another cell-division protein, DedD, was filamentous and DrpB was found to interact with multiple divisome-associated proteins [45].

RcsB positively regulates a promoter of *ftsZ*, and multicopy RcsF is a known suppressor of the temperature sensitive *ftsZ84* mutant [3]. Although DrpB did not rescue the *ftsZ(Ts)* phenotype [45], it seemed possible that the ability of DrpB to act as a weak suppressor of the *ftsEX* mutant is due to its role as an Rcs activator. To test this hypothesis, we asked whether RcsB was needed for the ability of DrpB to suppress the growth defect of Δ*ftsEX* cells in the absence of salt. We overexpressed DrpB from a plasmid-encoded arabinose-inducible promoter in 11*ftsEX* and 11*ftsEX* 11*rcsB* strains and assayed colony formation on LB medium with and without NaCl. As seen in Fig. 6A, multicopy DrpB can rescue the growth defect of *ftsEX* mutants even in cells deleted for *rcsB*, demonstrating that the suppression of Δ*ftsEX* is not via Rcs activation. We also asked whether activation of Rcs by DrpB required *ftsEX*, using the P*_rprA_*::mCherry reporter in a Δ*ftsEX* strain. We found that DrpB indeed induces Rcs in this strain (Fig. 6B). Therefore, the role of DrpB in cell growth in the absence of Δ*ftsEX* appears to be independent of Rcs activation and vice versa.

**Fig. 6:**
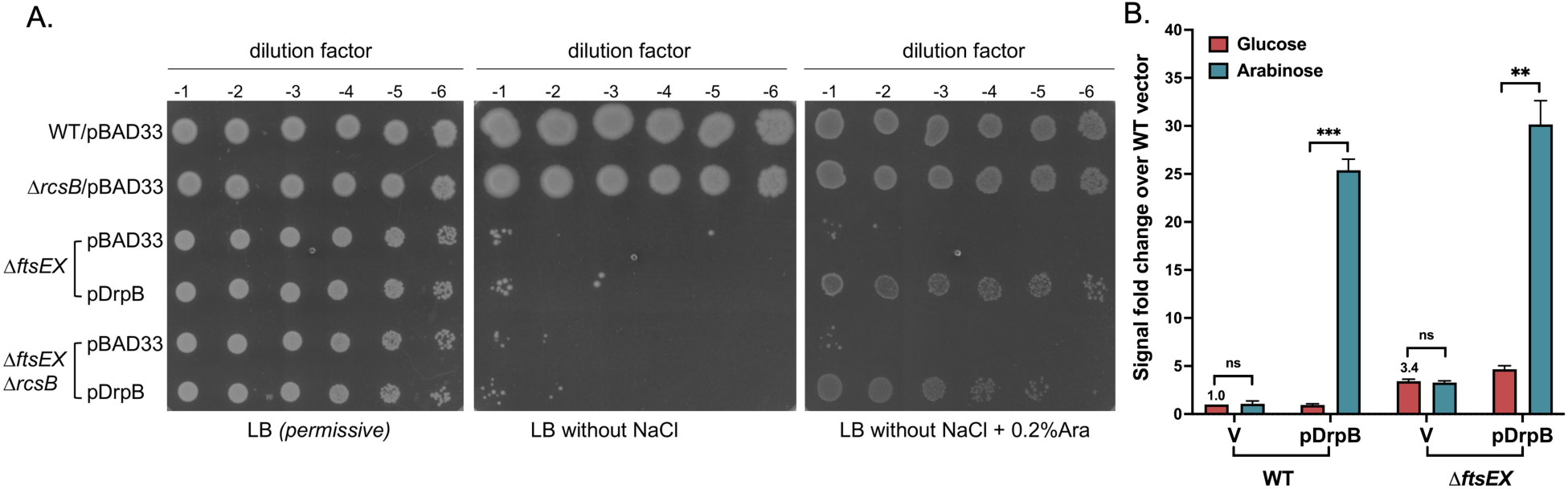
DrpB signaling is independent of *ftsEX*. **A. Role of DrpB as a *ftsEX* suppressor is independent of its role as an Rcs activator:** Strains transformed with the pBAD33 vector or pDrpB (pDSW1977) were grown overnight in LB Miller media with chloramphenicol at 37℃. The cultures were diluted to an OD_600_ of 1 and 4 ul dilutions were spotted on LB Miller (permissive), LB without NaCl, and LB with 0.2% arabinose with no NaCl. All plates contained chloramphenicol (25 μg/ml). Plates were imaged after 16h incubation at 37℃. The strains used are: WT (EC251), *rcsB::kan* (AP158, *ΔrcsB*), *ΔftsEX* (EC1215), and *ΔftsEX rcsB::kan* (AP159, *ΔftsEX ΔrcsB*). **B. DrpB can activate Rcs signaling in *ΔftsEX*:** For the P*_rprA_*::mCherry assay, the WT (EAW8) and *ftsE::kan* (AP154, *ΔftsEX*) strains transformed with the pBAD33 vector or pDrpB (pDSW1977) were grown in MOPS minimal glycerol medium (with 150 mM NaCl) containing chloramphenicol and either 0.2% glucose or 0.02% arabinose at 37℃. The RFU at OD 0.4 is plotted relative to uninduced WT with the vector, set to 1. Statistical significance was calculated using multiple unpaired *t*-tests and is shown as follows: ns (*P* > 0.05; non-significant),** (*P* ≤ 0.01), *** (*P* ≤ 0.001).

A set of DrpB mutants were constructed in the pBAD plasmid and tested both for Rcs induction and for ability to rescue the growth of Δ*ftsEX* in the absence of salt (Fig. S7). Since the periplasmic domain of RcsC is needed for DrpB signaling, it could be hypothesized that there is a periplasmic interaction between these proteins. Therefore, we constructed a DrpB mutant with a deletion of its periplasmic region (11peri_48-57_); this deletion had no effect on the ability of DrpB to activate Rcs (Fig. S7A). However, DrpB 11peri_48-57_ did not rescue the growth of the Δ*ftsEX* mutant, suggesting it is important for this function (Fig. S7B). An interaction between the transmembrane regions of DrpB and RcsC also seemed possible. Based on the sequence similarity with other DrpB homologs, we generated point mutants in four highly conserved amino acid residues in and around each of the TM helices of DrpB. Fig. S7A shows the effect of these mutants on Rcs signaling in a strain deleted for RcsF. Only DrpB T89A had lost the ability to activate Rcs. This allele only slightly decreased suppression of Δ*ftsEX* (Fig. S7B). The DrpB R38F had decreased but not absent activity for both induction of Rcs and suppression of Δ*ftsEX* (Fig. S7B). We cannot rule out that these mutants are less active due to mislocalization or decreased stability. However, overall the results confirm the existence of two separable functions of DrpB, with the periplasmic region critical for suppression of Δ*ftsEX* but not for Rcs induction, and T89A critical for Rcs induction but reasonably able to suppress 11*ftsEX*.

The BACTH assay was used to test if DrpB directly interacts with any of the Rcs components. However, full-length RcsC has previously been found to be non-functional in the BACTH assay [9], and was also negative for interaction with DrpB (Fig. S8A). We observed variable and weak (not significant) interactions of DrpB with both IgaA and RcsD (Fig. S8A). DrpB interacted with multiple divisome proteins as well as with MalF (used as a control) in a previous study, suggesting the possibility of false-positives in BACTH assays with other membrane proteins [45]

As with DjlA, we were not successful in defining an Rcs phenotype for deletion of *drpB*. The 11*drpB* strain responds to PMBN, A22 and mecillinam (Fig. S8B), which is consistent with a previous observation that DrpB is not needed for the Rcs response to osmotic shock or ethanol stress [15]. Multicopy DrpB also was previously found to induce the *psp* (phage shock) signaling pathway [15]. Rcs induction by DrpB was unaffected by the absence of *psp* components *pspC* and *pspF* (Fig. S8C), ruling out induction of Rcs indirectly via induction of Psp. More studies are needed to ascertain if DrpB activates the Rcs cascade by interacting directly with RcsC and the physiological conditions during which it functions as an Rcs activator.

Similar to DrpB, a group of small cytosolic proteins in the *ycgz-ymgABC* operon, which is involved in biofilm formation and acid-resistance, have been demonstrated to trigger the Rcs cascade [17, 49]. AriR/YmgB, in complex with YcgZ and YmgA, was reported to targts RcsC to induce the Rcs pathway [50]. We tested the Rcs activation by YmgB in our strains and found it to be RcsF independent and blocked by the RcsD T411A mutation (Fig. S9). However, in contrast with DrpB, it is independent of the periplasmic domain of RcsC, indicating a different mode of Rcs activation.

## Discussion

The Rcs phosphorelay is a remarkably sophisticated signaling pathway capable of responding to a wide array of stress signals at the cell envelope. RcsF plays a vital role in sensing most of the Rcs induction cues but is not strictly necessary for activating signaling. Our study elucidates diverse modes of activation of the Rcs phosphorelay which are independent of RcsF (Fig 7). The results demonstrate that RcsF is not the sole sensor for this pathway. In particular, our identification of DrpB as an RcsF-independent activator suggests the possibility of other yet unidentified RcsF-independent activators and particularly activators that act via the RcsC periplasmic domain.

**Fig. 7:**
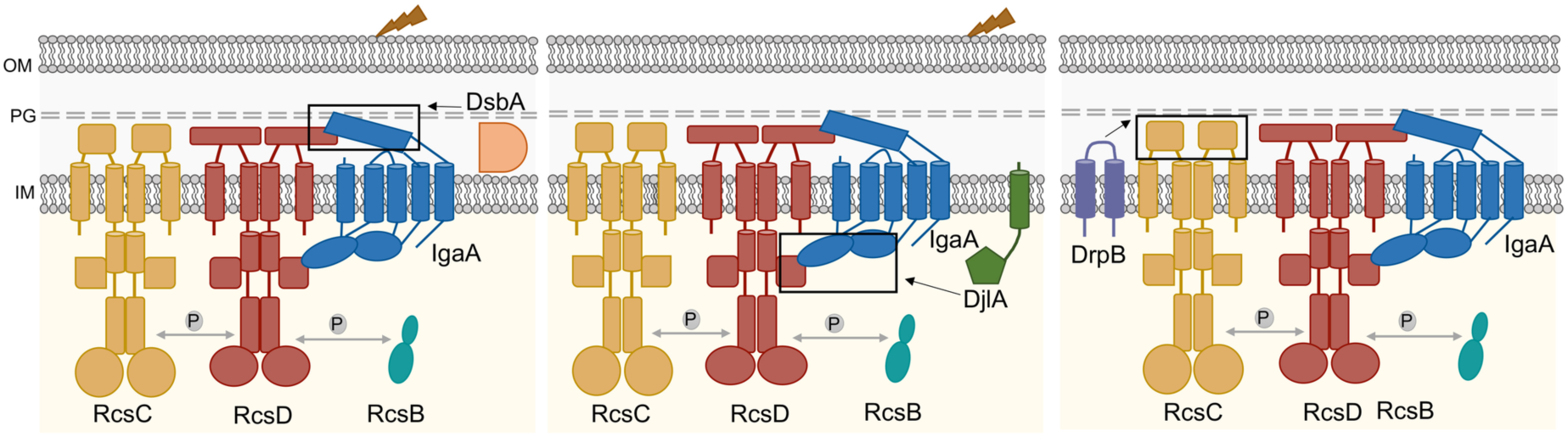
Diverse modes of RcsF-independent activation by DsbA, DjlA, and DrpB. First, *dsbA* mutants activate Rcs signaling in response to DTT and PMBN. The absence of DsbA likely leads to defective disulfide bond formation and misfolding of the IgaA periplasmic domain. Thus, DsbA is needed to maintain proper IgaA-RcsD interactions in the periplasm and in its absence, IgaA cannot effectively regulate Rcs signaling. Second, DjlA acts as a cochaperone for IgaA-RcsD interactions. Overexpression of DjlA weakens the IgaA-RcsD interactions in the cytoplasm leading to Rcs activation. Third, DrpB overexpression induces the Rcs cascade but it does not act directly on IgaA-RcsD interactions like DsbA and DjlA. Uniquely, DrpB requires the RcsC periplasmic domain to activate Rcs signaling and either activates it by direct interaction with RcsC or via an indirect mechanism involving RcsC as the sensor instead of RcsF.

We delineate how three RcsF-independent activators trigger the Rcs phosphorelay, each in a different fashion. Our current understanding of the regulation of Rcs signaling is primarily based on the interactions of IgaA with RcsF and RcsD. Interactions of RcsF with IgaA form the first level of regulation crucial for activation of signaling (Fig. 1A). Signals detected by RcsF change the interaction of IgaA with RcsD; this IgaA-RcsD interaction serves as the regulatory switch necessary for signal activation. Dependent on the nature of these interactions, the RcsC histidine kinase acts as either a phosphatase (RcsB response regulator not phosphorylated, inactive) or a kinase (RcsB response regulator phosphorylated and active). The IgaA-RcsD interactions are also essential for repression of signaling, thereby preventing constitutive activation and the resulting lethality. Our work underscores the importance of these IgaA-RcsD interactions in regulating this cascade even in the absence of RcsF-dependent activation. Signaling in all three cases, in cells deleted for DsbA or overproducing DjlA or DrpB, is dependent on RcsC and RcsD and does not initiate signaling merely by increasing expression of components such as RcsB.

### Interrupting the interactions of RcsD and IgaA induce Rcs signalling

Loss of DsbA or DjlA overproduction each activate signaling by weakening the interactions between IgaA and RcsD, affecting their folding/conformation via distinct modes. We have previously shown that IgaA and RcsD interact both through their periplasmic domains and their cytoplasmic domains ([9], Fig. 1A). The periplasmic strong contact provides an anchor that keeps signaling from rising too high. Based on the identification of the RcsDT411A mutant in the cytoplasmic PAS domain of RcsD that blocks RcsF-dependent induction, the cytoplasmic contact was defined as the regulatory switch region. The loss of DsbA impacts the periplasmic contacts, likely due to its role in formation of disulfide bonds in IgaA, and therefore IgaA folding. This is reflected in a decrease in the interaction of IgaA and RcsD in the absence of DsbA and in elevated Rcs signaling, further increased when cells are treated with DTT. Presumably in a strain devoid of DsbA, a fraction of IgaA is in a reduced state with one or both disulfide bonds disrupted; addition of DTT leads to the disruption of any remaining disulfide bonds. Mutating the IgaA periplasmic cysteines predicted to be involved in these disulfide bonds mimics the *dsbA* mutant, decreasing the interaction of IgaA and RcsD (Fig. S4C) and increasing signaling. RcsF itself requires disulfide bonds for proper folding and function [30, 31]. Therefore, it is not surprising that *dsbA* mutants no longer recognize RcsF-dependent cues such as A22 and mecillinam (Fig. S4A), since RcsF is non-functional in this strain. Similar observations have been made in *Salmonella*; *Salmonella dsbA* mutants activate Rcs, independent of RcsF, and this is increased upon DTT addition, while a strain carrying a *bamB* deletion induces Rcs, dependent upon RcsF and is unaffected by DTT treatment [51].

We might have also expected PMBN, which normally induces the Rcs system in an RcsF-dependent fashion (see Fig. 1B) to be ineffective in the *dsbA* mutant. However, it induces in much the manner that DTT does, independently of *rcsF* (Fig. 1B). This result suggests that PMBN causes some redox stress, in addition to the LPS/membrane damage that leads to RcsF-dependent induction. When the Dsb network is functional, DTT cannot induce Rcs signaling and any PMBN induction requires RcsF, suggesting that Rcs probably does not monitor disulfide bond formation in the periplasm as part of its normal function. Instead, the disulfide bonds are necessary for proper folding of IgaA to allow the periplasmic interaction with RcsD. We cannot rule out the possibility that under extreme oxidative stress, IgaA disulfide bonds might be disrupted sufficiently to induce Rcs signaling. A study of the roles of the periplasmic cysteines of IgaA in *Salmonella* also found that disulfide bonds were critical both for proper repression of Rcs and, to some extent, for viability of the cells [33].

Overproduction of DjlA, a membrane-localized DnaJ family protein, also loosens the interaction of IgaA and RcsD (Fig. 5A) and turns up signaling independently of RcsF (Fig. 1B); these effects depend on an active J domain (mutated in DjlA H233Q). However, in this case, the primary effect is likely on the cytoplasmic domains of IgaA, RcsD, or both of these proteins. This is suggested both by the observation that DjlA overproduction but not loss of DsbA overcomes the RcsD T411A mutation (Fig. 3) and by the location of the J domain in the cytoplasm. The ability of DjlA overproduction to induce signaling also allows us to conclude that the interaction of the RcsD PAS-like domain with the IgaA cytoplasmic domain can regulate activation by both RcsF-dependent and RcsF-independent signals.

DjlA was the only RcsF-independent activator which overcame the heightened interaction between RcsD T411A mutant and IgaA. This would be consistent with DjlA acting as a chaperone to mediate these interactions. Molecular chaperone systems can assist protein-protein interactions by loosening and altering protein conformations. Unlike DnaJ and CbpA, the other *E. coli* DnaJ family proteins capable of working with DnaK, DjlA does not carry a specific substrate-binding domain, so it is not known how it identifies its clients [42]. Since only DjlA, and not the other two DnaJ family proteins, induces Rcs, it seemed possible that the membrane insertion of DjlA is critical in recruiting DnaK to the sites of IgaA/RcsD. The transmembrane helix of DjlA contains several conserved residues and it has been postulated earlier that this helix helps in substrate interaction [42]. This notion is reinforced by the lack of signaling from a cytosolic soluble form of DjlA ([13, 42], Fig. S5B). The transmembrane helix is also a dimerization helix, and DjlA fused to any other transmembrane helix was previously found to be incapable of activating this system [44]. However, we found that DjlA fused to the MalF transmembrane domains can activate Rcs, strongly suggesting this helix is not required for Rcs interaction. Our results also rule out RcsC as the DjlA substrate for Rcs activation. Either IgaA or RcsD, or the complex could be a target of DjlA; however, the ability of DjlA to disrupt interaction of IgaA11cyt and RcsD (Fig. S5C) is most consistent with RcsD as the target. Because the periplasmic interaction of RcsD and IgaA is sufficient for a strong interaction signal in the BACTH assay, we can also conclude that the action of DjlA on the cytoplasmic domain of RcsD must lead to a change in the RcsD periplasmic domain.

The effects here were seen when DjlA was overproduced. We do not know whether or when the chromosomally encoded DjlA is able to modulate the IgaA-RcsD interaction. Deletion of *djlA* strengthened the interaction of IgaA and RcsD in the BACTH assay (Fig. S6D) but did not have any effect on Rcs induction under a variety of conditions (Fig. S6). Not much is known about the physiological signals regulating DjlA expression; DjlA-stimulation of Rcs might be relevant under specific growth conditions or might be important for particular inducing signals not tested here. There is also evidence that DjlA acts as a negative regulator of Rcs [5, 52]. Those studies demonstrated a stimulation of the Rcs signal upon deletion of DjlA, particularly in mutant strains such as *mdo* and *rfa* which are already activated for Rcs in an RcsF-dependent manner. Lack of DjlA could increase the envelope stress caused by mutants in *mdo* and *rfa* independently of the RcsF-independent effect seen here. Undoubtedly, DjlA has substrates other than the Rcs system, and possibly the opposing effects (controlling envelope stress to negatively regulate Rcs and overcoming the RcsD/IgaA protein to induce Rcs) balance each other, making it more difficult to define a single copy effect.

### Rcs Activation via Histidine Kinase RcsC

Uniquely among the three proteins we investigated, DrpB required the RcsC periplasmic domain for signaling. RcsC is a member of a broad family of histidine kinases; in many of these, the periplasmic domains are critical for sensing environmental signals and regulating histidine kinase activity [53-55]. In our previous work, we found that IgaA interacted with RcsD, the phosphorelay protein, rather than RcsC and Rcs inducing treatments were independent of the RcsC periplasmic domain [9]. Thus, while RcsC activity (kinase vs. phosphatase) must be adjusted by RcsF-dependent induction, there was no evidence for a role of RscC in signal sensing. Thus, the identification of DrpB and its dependence on the RcsC periplasmic region demonstrates the likelihood that some signals act in the “classical” way seen for other histidine kinases; if so, such signals would be independent of RcsF. While we do not know if DrpB acts directly or indirectly on RcsC, it is tempting to speculate that the RcsC periplasmic domain is sensing a signal created byDrpB overexpression, allowing RcsC to activate signaling. Interestingly, a ribosome profiling study revealed DrpB as one of the proteins upregulated during severe acid stress in *E. coli* and this may be linked to its Rcs function [56]. Alternatively, the overproduction of DrpB itself is a source of stress which is detected by RcsC. DrpB overproduction also leads to the induction of another envelope stress response, the Psp response pathway [15]. The Psp response primarily detects and mitigates stress at the inner membrane (reviewed in [57]). While we find that the Psp pathway is not necessary for Rcs induction by DrpB (Fig. S8), it will be of interest to see whether other inducing signals, shared by these two systems, also share dependence upon the RcsC periplasmic domain.

Further evidence for RcsC as a sensor comes from studies on the protein YmgB. YmgB is a cytoplasmic protein involved in biofilm formation and acid-resistance and it stimulates the Rcs pathway, reportedly by interacting with the RcsC cytoplasmic domain [49, 50]. In our hands, and consistent with the model suggested by DrpB overexpression, YmgB-dependent induction is RcsF-independent, but, unlike DrpB, is also independent of the periplasmic domain of RcsC (Fig. S9). This is consistent with previous work on YmgB and raises the possibility that RcsC in fact senses different signals with its cytoplasmic and periplasmic domains.

A number of other proteins have been found to affect Rcs activity. YpdI, a lipoprotein of unknown function, induces mucoidy in the absence of RcsF [16]. Activation of Rcs in a double mutant of *yqjA* and *yghB*, both membrane proteins and putative lipid flippases, is partially dependent on RcsF [20, 58]. Deciphering their mechanisms could add to our knowledge about Rcs regulation.

Overall, our results emphasize the versatility and flexibility of the Rcs pathway. RcsF senses stress at the outer cell surface and in the periplasm. The RcsF-independent signaling pathways studied here suggest that, in addition, RcsC may play a role in sensing stressors at the cytoplasmic membrane, and possibly in the cytoplasm as well. Possibly this sensing by RcsC represents the remnants of the original version of this system, before RcsD, IgaA and RcsF evolved to sense a broader set of inducing signals. Clearly there is still much to be learned about this system, across species, and in different natural environments. Structural information on the Rcs proteins and how they interact is also still lacking. We look forward to better understanding this unique system in the future.

## Materials and Methods

### Bacterial growth conditions and strain construction

The strains were grown in LB medium with appropriate antibiotics (ampicillin 100 μg/ml, kanamycin 30-50 μg/ml, chloramphenicol 10 μg/ml for *cat-sacB* allele or 25 μg/ml for chl^R^ plasmids, tetracycline 25μg/ml, zeocin 50μg/ml). Glucose at a final concentration of 1% was added to reduce the basal expression from the plasmids containing pBAD and pLac promoters.

Strains, plasmids, primers and synthetic DNA fragments (gBlocks) used in this study are listed in Tables S1, S2, S3, and S4 respectively. Oligonucleotides and gBlocks were from IDT DNA, Coralville, IA. Strains were constructed by recombineering or P1 transduction with selectable markers (Table S1). Recombineering was done in strains carrying a chromosomal mini-λ Red system (*miniλ::tet*) or a plasmid-borne Red system (pSIM27). Some strains were generated by direct P1 transduction from the corresponding mutant strains in the Keio collection [59]. Plasmids were constructed by the Gibson assembly method using the In-fusion HD Cloning kit (Takara Bio USA) [60]. PCR products were purified using column purification (Qiagen) and transformed into NEB DH5α F’*lacIQ* cells. Site-directed mutagenesis in the genes was carried out using the QuikChange Site-directed mutagenesis kit (Agilent). Sequencing was done to confirm the chromosomal and plasmid modifications.

### Screening for multicopy RcsF-independent activators

A pBR-based plasmid library [61] was transformed into EAW34 electrocompetent cells. Aliquots of the transformed mixture were plated on LB plates containing 100 μg/ml ampicillin and incubated overnight at 37°C. Subsequently, the plates were imaged and the colonies displaying high fluorescence were restreaked on LB ampicillin plates and purified. The plasmid DNA was isolated from these colonies and retransformed into EAW34 to validate their phenotype. The positive candidate plasmids were isolated, sequenced and the reads mapped to the *E. coli K-12 MG1655* genome to identify the inserts.

### Fluorescence reporter assays

Growth and fluorescence assays for Rcs activation were carried out in 96-well plates in a Tecan Spark microplate reader. All the strains carried a P*_rprA_*::mCherry transcriptional fusion at the *ara* locus as a reporter for Rcs signaling [9]. For strains carrying overexpression plasmids expressing DjlA, DrpB, or RcsF, the overnight cultures were grown in MOPS minimal glucose (0.2%) medium (Teknova). For the P*_rprA_*::mCherry assays for these strains, they were diluted to OD_600_ 0.03-0.05 in MOPS minimal glycerol media (0.05% glucose, 0.5% glycerol) with 0.02% arabinose (induced) or 0.2% glucose as an uninduced control. Both the optical density (OD_600_) and mCherry fluorescence were monitored every fifteen minutes for 12 hours. Polymyxin B nonapeptide (PMBN; Sigma) was used at 20 μg/ml, to induce the Rcs system. A22 (an MreB inhibitor) was used at 5 μg/ml, and mecillinam was used at 0.3 μg/ml. Each assay was performed in technical duplicates in the microtiter plate, with the biological replicates performed on different days. The fluorescence values at equivalent OD_600_ values (0.4 ± 0.04) for each strain were converted to bar graphs and plotted as signal fold change of fluorescence signal over the uninduced wild type strain. Error bars represent the standard deviation of at least 3 assays. For strains that did not reach an OD_600_ value of 0.4, the OD is noted on the bar graph. For the P*_rprA_*::mCherry assays for the strains not carrying any plasmids, MOPS minimal glucose media was used for growing overnight cultures and performing the fluorescence assays. For the assays pertaining to *dsbA* mutants, the overnight cultures were grown in LB medium. Subsequently, cells were washed with and diluted to OD_600_ 0.05 in MOPS minimal glucose media to carry out the P*_rprA_*::mCherry assay. To activate the Rcs system, Dithiothreitol (DTT, Sigma) was added at a final concentration of 1mM or Polymyxin B nonapeptide (PMBN; Sigma), was used at 20 μg/ml. All assays were performed at 37°C, unless otherwise indicated. Values derived from at least three independent experiments were plotted to show the mean with error bars indicating standard deviation. These were statistically analyzed by multiple unpaired *t*-tests using GraphPad Prism 10 software.

### Bacterial adenylate cyclase two hybrid assay (BACTH)

For the bacterial adenylate cyclase two hybrid assay (BACTH), an adenylate cyclase mutant strain (BTH101) or a derivative if this strain was used. The proteins whose interactions were being examined were cloned into the T18 and T25 portions of adenylate cyclase [34, 35]. The reconstitution of adenylate cyclase upon their interaction allows production of cAMP, which interacts with CRP to activate the *lac* operon, assayed as beta-galactosidase activity. In the absence of interaction, T18 and T25 do not get reconstituted to form a functional adenylate cyclase and the strain does not express beta-galactosidase. The proteins used in this study were fused at their C-terminal to the Cya fragments. Plasmids expressing IgaA or RcsD as T18/T25 fusion constructs were cotransformed into BTH101, plated on LB-agar medium containing 100 μg/ml ampicillin and 50 μg/ml kanamycin and incubated at 30°C for 2 days. To measure the beta-galactosidase activity, the resulting colonies were inoculated and grown overnight in LB medium containing 100 μg/ml ampicillin, 50 μg/ml kanamycin, and 0.5 mM IPTG at 30°C. The beta-galactosidase assay was performed in 96-well plates. The activity was analysed by measuring the kinetics of ONPG degradation by monitoring the OD 420 nm at 28°C for 40 min at intervals of 1 min in a Tecan Spark microplate reader. The beta-galactosidase activity was calculated using the slope of OD 420 divided by their OD 600. Each protein fusion paired with the cognate vector produces very little activity and these were used as negative controls. Every graph is compiled from at least 3 separate sets of assays, and the beta-galactosidase activity is plotted relative to the IgaA/RcsD interaction in that particular experiment.

To test the effect of DsbA activity on the various IgaA/RcsD constructs, the BACTH assays were carried out in BTH101 and a *dsbA* mutant derivative of BTH101 (AP58). To test the effect of DjlA on the IgaA/RcsD interactions, d*jlA* or mutants of *djlA* were cloned downstream of IgaA under the control of the same promoter. The assay was carried out in BTH101 or its derivatives EAW1, EAW2, and EAW4 as above and interactions plotted relative to the IgaA/RcsD interaction in WT, set to 1.

### Western blots

For Fig. S4D, plasmids were transformed into EAW8 and EAW62 and for Fig. S5D, the plasmids were transformed into *E. coli* DH5α competent cells (NEB). The colonies were inoculated into LB media containing Amp (100 μg/ml) and 1 mM IPTG and then grown for 8 hours at 30°C. One ml of culture was precipitated using TCA, washed in acetone and resuspended in LDS sample buffer standardized to OD. 8 μl volumes were loaded on 4–12% NuPage gradient gels (Invitrogen, CA) and then transferred onto nitrocellulose membranes in an iBlot2 transfer device as per manufacturer’s specifications (Invitrogen, CA). The membranes were blocked with Casein PBS and then incubated in CyaA-T18 mouse monoclonal primary antibody at 1:10,000 (Santa Cruz Biotechnology, CA) or/and a mouse monoclonal anti-EF-Tu at 1:10,000 (Hycult Biotech, PA) overnight at 4°C. Secondary fluorescent antibody anti-mouse DyLight800 (Bio-Rad, CA) was used at 1:10,000. Imaging was done on a ChemiDoc MP imaging system (Bio-Rad).

### Viability testing of *ftsEX* mutants

The wildtype and *ftsEX* deletion strains were transformed with the pBAD33 vector or a DrpB containing plasmid and plated on LB Miller-agar medium at 30°C containing 25 μg/ml chloramphenicol. Overnight cultures were grown in LB Miller medium 25 μg/ml chloramphenicol at 30°C. Cells were harvested from 1 mL of culture and diluted to OD_600_ 1.0 in LB Miller or LB0N (LB without any NaCl) medium. Serial dilutions (10-fold) were made and 4 μl was spotted onto LB Miller-agar or LB0N-agar plates containing 25 μg/ml chloramphenicol. Arabinose was added at 0.2% for induction in LB0N. Plates were imaged after 16 h of growth at 37°C.

## Supporting information

Petchiappan et al Supplemental Figs. and Tables

## Acknowledgements

This research was supported by the Intramural Research Program of the NIH, National Cancer Institute, Center for Cancer Research. We thank members of the Gottesman lab for comments on the manuscript. EAW was supported by a Fi2 PRAT fellowship award from NIGMS. Tae Yoon Chang, a summer student, performed the initial library screen for RcsF-independent Rcs activation that identified DrpB as an activator. We thank David Weiss for discussions and for providing strains (see Table S1).

## References

1. Silhavy TJ, Kahne D, Walker S. The bacterial cell envelope. Cold Spring Harb Perspect Biol. 2010;2(5):a000414. Epub 20100414. doi: 10.1101/cshperspect.a000414. PubMed PMID: 20452953; PubMed Central PMCID: PMCPMC2857177.

2. Wall E, Majdalani N, Gottesman S. The Complex Rcs Regulatory Cascade. Annu Rev Microbiol. 2018;72:111–39. Epub 20180613. doi: 10.1146/annurev-micro-090817-062640. PubMed PMID: 29897834.

3. Gervais FG, Drapeau GR. Identification, cloning, and characterization of rcsF, a new regulator gene for exopolysaccharide synthesis that suppresses the division mutation ftsZ84 in Escherichia coli K-12. J Bacteriol. 1992;174(24):8016–22. doi: 10.1128/jb.174.24.8016-8022.1992. PubMed PMID: 1459951; PubMed Central PMCID: PMCPMC207539.

4. Majdalani N, Gottesman S. The Rcs phosphorelay: a complex signal transduction system. Annu Rev Microbiol. 2005;59:379–405. doi: 10.1146/annurev.micro.59.050405.101230. PubMed PMID: 16153174.

5. Castanié-Cornet MP, Cam K, Jacq A. RcsF is an outer membrane lipoprotein involved in the RcsCDB phosphorelay signaling pathway in Escherichia coli. J Bacteriol. 2006;188(12):4264–70. doi: 10.1128/jb.00004-06. PubMed PMID: 16740933; PubMed Central PMCID: PMCPMC1482940.

6. Laubacher ME, Ades SE. The Rcs phosphorelay is a cell envelope stress response activated by peptidoglycan stress and contributes to intrinsic antibiotic resistance. J Bacteriol. 2008;190(6):2065–74. Epub 20080111. doi: 10.1128/jb.01740-07. PubMed PMID: 18192383; PubMed Central PMCID: PMCPMC2258881.

7. Farris C, Sanowar S, Bader MW, Pfuetzner R, Miller SI. Antimicrobial peptides activate the Rcs regulon through the outer membrane lipoprotein RcsF. J Bacteriol. 2010;192(19):4894–903. Epub 20100730. doi: 10.1128/jb.00505-10. PubMed PMID: 20675476; PubMed Central PMCID: PMCPMC2944553.

8. Cano DA, Domínguez-Bernal G, Tierrez A, Garcia-Del Portillo F, Casadesús J. Regulation of capsule synthesis and cell motility in Salmonella enterica by the essential gene igaA. Genetics. 2002;162(4):1513–23. doi: 10.1093/genetics/162.4.1513. PubMed PMID: 12524328; PubMed Central PMCID: PMCPMC1462382.

9. Wall EA, Majdalani N, Gottesman S. IgaA negatively regulates the Rcs Phosphorelay via contact with the RcsD Phosphotransfer Protein. PLoS Genet. 2020;16(7):e1008610. Epub 20200727. doi: 10.1371/journal.pgen.1008610. PubMed PMID: 32716926; PubMed Central PMCID: PMCPMC7418988.

10. Cho SH, Szewczyk J, Pesavento C, Zietek M, Banzhaf M, Roszczenko P, et al. Detecting envelope stress by monitoring β-barrel assembly. Cell. 2014;159(7):1652–64. doi: 10.1016/j.cell.2014.11.045. PubMed PMID: 25525882.

11. Konovalova A, Mitchell AM, Silhavy TJ. A lipoprotein/β-barrel complex monitors lipopolysaccharide integrity transducing information across the outer membrane. Elife. 2016;5. Epub 20160610. doi: 10.7554/eLife.15276. PubMed PMID: 27282389; PubMed Central PMCID: PMCPMC4942254.

12. Zuber M, Hoover TA, Court DL. Analysis of a Coxiella burnetti gene product that activates capsule synthesis in Escherichia coli: requirement for the heat shock chaperone DnaK and the two-component regulator RcsC. J Bacteriol. 1995;177(15):4238–44. doi: 10.1128/jb.177.15.4238-4244.1995. PubMed PMID: 7635811; PubMed Central PMCID: PMCPMC177168.

13. Kelley WL, Georgopoulos C. Positive control of the two-component RcsC/B signal transduction network by DjlA: a member of the DnaJ family of molecular chaperones in Escherichia coli. Mol Microbiol. 1997;25(5):913–31. doi: 10.1111/j.1365-2958.1997.mmi527.x. PubMed PMID: 9364917.

14. Clavel T, Lazzaroni JC, Vianney A, Portalier R. Expression of the tolQRA genes of Escherichia coli K-12 is controlled by the RcsC sensor protein involved in capsule synthesis. Mol Microbiol. 1996;19(1):19–25. doi: 10.1046/j.1365-2958.1996.343880.x. PubMed PMID: 8821933.

15. Bury-Moné S, Nomane Y, Reymond N, Barbet R, Jacquet E, Imbeaud S, et al. Global analysis of extracytoplasmic stress signaling in Escherichia coli. PLoS Genet. 2009;5(9):e1000651. Epub 20090918. doi: 10.1371/journal.pgen.1000651. PubMed PMID: 19763168; PubMed Central PMCID: PMCPMC2731931.

16. Potrykus J, Wegrzyn G. The ypdI gene codes for a putative lipoprotein involved in the synthesis of colanic acid in Escherichia coli. FEMS Microbiol Lett. 2004;235(2):265–71. doi: 10.1016/j.femsle.2004.04.041. PubMed PMID: 15183873.

17. Tschowri N, Busse S, Hengge R. The BLUF-EAL protein YcgF acts as a direct anti-repressor in a blue-light response of Escherichia coli. Genes Dev. 2009;23(4):522–34. doi: 10.1101/gad.499409. PubMed PMID: 19240136; PubMed Central PMCID: PMCPMC2648647.

18. Westphal K, Langklotz S, Thomanek N, Narberhaus F. A trapping approach reveals novel substrates and physiological functions of the essential protease FtsH in Escherichia coli. J Biol Chem. 2012;287(51):42962–71. Epub 20121022. doi: 10.1074/jbc.M112.388470. PubMed PMID: 23091052; PubMed Central PMCID: PMCPMC3522291.

19. Bittner LM, Westphal K, Narberhaus F. Conditional Proteolysis of the Membrane Protein YfgM by the FtsH Protease Depends on a Novel N-terminal Degron. J Biol Chem. 2015;290(31):19367–78. Epub 20150619. doi: 10.1074/jbc.M115.648550. PubMed PMID: 26092727; PubMed Central PMCID: PMCPMC4521054.

20. Sikdar R, Simmons AR, Doerrler WT. Multiple envelope stress response pathways are activated in an Escherichia coli strain with mutations in two members of the DedA membrane protein family. J Bacteriol. 2013;195(1):12–24. Epub 20121005. doi: 10.1128/jb.00762-12. PubMed PMID: 23042993; PubMed Central PMCID: PMCPMC3536178.

21. Majdalani N, Hernandez D, Gottesman S. Regulation and mode of action of the second small RNA activator of RpoS translation, RprA. Mol Microbiol. 2002;46(3):813–26. doi: 10.1046/j.1365-2958.2002.03203.x. PubMed PMID: 12410838.

22. Brill JA, Quinlan-Walshe C, Gottesman S. Fine-structure mapping and identification of two regulators of capsule synthesis in Escherichia coli K-12. J Bacteriol. 1988;170(6):2599–611. doi: 10.1128/jb.170.6.2599-2611.1988. PubMed PMID: 2836365; PubMed Central PMCID: PMCPMC211177.

23. Shiba Y, Miyagawa H, Nagahama H, Matsumoto K, Kondo D, Matsuoka S, et al. Exploring the relationship between lipoprotein mislocalization and activation of the Rcs signal transduction system in Escherichia coli. Microbiology (Reading). 2012;158(Pt 5):1238–48. Epub 20120209. doi: 10.1099/mic.0.056945-0. PubMed PMID: 22322964.

24. Heras B, Shouldice SR, Totsika M, Scanlon MJ, Schembri MA, Martin JL. DSB proteins and bacterial pathogenicity. Nat Rev Microbiol. 2009;7(3):215-25. Epub 20090209. doi: 10.1038/nrmicro2087. PubMed PMID: 19198617.

25. Bardwell JC, McGovern K, Beckwith J. Identification of a protein required for disulfide bond formation in vivo. Cell. 1991;67(3):581–9. doi: 10.1016/0092-8674(91)90532-4. PubMed PMID: 1934062.

26. Kadokura H, Beckwith J. Four cysteines of the membrane protein DsbB act in concert to oxidize its substrate DsbA. Embo j. 2002;21(10):2354–63. doi: 10.1093/emboj/21.10.2354. PubMed PMID: 12006488; PubMed Central PMCID: PMCPMC126001.

27. Berkmen M, Boyd D, Beckwith J. The nonconsecutive disulfide bond of Escherichia coli phytase (AppA) renders it dependent on the protein-disulfide isomerase, DsbC. J Biol Chem. 2005;280(12):11387–94. Epub 20050110. doi: 10.1074/jbc.M411774200. PubMed PMID: 15642731.

28. Missiakas D, Georgopoulos C, Raina S. Identification and characterization of the Escherichia coli gene dsbB, whose product is involved in the formation of disulfide bonds in vivo. Proc Natl Acad Sci U S A. 1993;90(15):7084–8. doi: 10.1073/pnas.90.15.7084. PubMed PMID: 7688471; PubMed Central PMCID: PMCPMC47080.

29. Dailey FE, Berg HC. Mutants in disulfide bond formation that disrupt flagellar assembly in Escherichia coli. Proc Natl Acad Sci U S A. 1993;90(3):1043–7. doi: 10.1073/pnas.90.3.1043. PubMed PMID: 8503954; PubMed Central PMCID: PMCPMC45807.

30. Kadokura H, Tian H, Zander T, Bardwell JC, Beckwith J. Snapshots of DsbA in action: detection of proteins in the process of oxidative folding. Science. 2004;303(5657):534-7. doi: 10.1126/science.1091724. PubMed PMID: 14739460.

31. Leverrier P, Declercq JP, Denoncin K, Vertommen D, Hiniker A, Cho SH, Collet JF. Crystal structure of the outer membrane protein RcsF, a new substrate for the periplasmic protein-disulfide isomerase DsbC. J Biol Chem. 2011;286(19):16734–42. Epub 20110316. doi: 10.1074/jbc.M111.224865. PubMed PMID: 21454485; PubMed Central PMCID: PMCPMC3089515.

32. Rogov VV, Rogova NY, Bernhard F, Löhr F, Dötsch V. A disulfide bridge network within the soluble periplasmic domain determines structure and function of the outer membrane protein RCSF. J Biol Chem. 2011;286(21):18775–83. Epub 20110406. doi: 10.1074/jbc.M111.230185. PubMed PMID: 21471196; PubMed Central PMCID: PMCPMC3099694.

33. Pucciarelli MG, Rodríguez L, García-Del Portillo F. A Disulfide Bond in the Membrane Protein IgaA Is Essential for Repression of the RcsCDB System. Front Microbiol. 2017;8:2605. Epub 20171222. doi: 10.3389/fmicb.2017.02605. PubMed PMID: 29312270; PubMed Central PMCID: PMCPMC5744062.

34. Battesti A, Bouveret E. The bacterial two-hybrid system based on adenylate cyclase reconstitution in Escherichia coli. Methods. 2012;58(4):325–34. Epub 20120724. doi: 10.1016/j.ymeth.2012.07.018. PubMed PMID: 22841567.

35. Karimova G, Pidoux J, Ullmann A, Ladant D. A bacterial two-hybrid system based on a reconstituted signal transduction pathway. Proc Natl Acad Sci U S A. 1998;95(10):5752–6. doi: 10.1073/pnas.95.10.5752. PubMed PMID: 9576956; PubMed Central PMCID: PMCPMC20451.

36. Watanabe N, Savchenko A. Molecular insights into the initiation step of the Rcs signaling pathway. Structure. 2024. Epub 20240625. doi: 10.1016/j.str.2024.06.003. PubMed PMID: 38964336.

37. Clarke DJ, Jacq A, Holland IB. A novel DnaJ-like protein in Escherichia coli inserts into the cytoplasmic membrane with a type III topology. Mol Microbiol. 1996;20(6):1273–86. doi: 10.1111/j.1365-2958.1996.tb02646.x. PubMed PMID: 8809778.

38. Alekhina O, Valkovicova L, Turna J. Study of membrane attachment and in vivo co-localization of TerB protein from uropathogenic Escherichia coli KL53. Gen Physiol Biophys. 2011;30(3):286–92. doi: 10.4149/gpb_2011_03_286. PubMed PMID: 21952438.

39. Anantharaman V, Iyer LM, Aravind L. Ter-dependent stress response systems: novel pathways related to metal sensing, production of a nucleoside-like metabolite, and DNA-processing. Mol Biosyst. 2012;8(12):3142–65. doi: 10.1039/c2mb25239b. PubMed PMID: 23044854; PubMed Central PMCID: PMCPMC4104200.

40. Genevaux P, Wawrzynow A, Zylicz M, Georgopoulos C, Kelley WL. DjlA is a third DnaK co-chaperone of Escherichia coli, and DjlA-mediated induction of colanic acid capsule requires DjlA-DnaK interaction. J Biol Chem. 2001;276(11):7906–12. Epub 20001205. doi: 10.1074/jbc.M003855200. PubMed PMID: 11106641.

41. Bernard S, Clarke DJ, Chen MX, Holland IB, Jacq A. Increased sensitivity of E. coli to novobiocin, EDTA and the anticalmodulin drug W7 following overproduction of DjlA requires a functional transmembrane domain. Mol Gen Genet. 1998;259(6):645-55. doi: 10.1007/pl00008629. PubMed PMID: 9819058.

42. Clarke DJ, Holland IB, Jacq A. Point mutations in the transmembrane domain of DjlA, a membrane-linked DnaJ-like protein, abolish its function in promoting colanic acid production via the Rcs signal transduction pathway. Mol Microbiol. 1997;25(5):933–44. doi: 10.1111/j.1365-2958.1997.mmi528.x. PubMed PMID: 9364918.

43. Genevaux P, Schwager F, Georgopoulos C, Kelley WL. The djlA gene acts synergistically with dnaJ in promoting Escherichia coli growth. J Bacteriol. 2001;183(19):5747–50. doi: 10.1128/jb.183.19.5747-5750.2001. PubMed PMID: 11544239; PubMed Central PMCID: PMCPMC95468.

44. Toutain CM, Clarke DJ, Leeds JA, Kuhn J, Beckwith J, Holland IB, Jacq A. The transmembrane domain of the DnaJ-like protein DjlA is a dimerisation domain. Mol Genet Genomics. 2003;268(6):761–70. Epub 20030131. doi: 10.1007/s00438-002-0793-z. PubMed PMID: 12655402.

45. Yahashiri A, Babor JT, Anwar AL, Bezy RP, Piette EW, Arends SJR, et al. DrpB (YedR) Is a Nonessential Cell Division Protein in Escherichia coli. J Bacteriol. 2020;202(23). Epub 20201104. doi: 10.1128/jb.00284-20. PubMed PMID: 32900831; PubMed Central PMCID: PMCPMC7648144.

46. Schmidt KL, Peterson ND, Kustusch RJ, Wissel MC, Graham B, Phillips GJ, Weiss DS. A predicted ABC transporter, FtsEX, is needed for cell division in Escherichia coli. J Bacteriol. 2004;186(3):785–93. doi: 10.1128/jb.186.3.785-793.2004. PubMed PMID: 14729705; PubMed Central PMCID: PMCPMC321481.

47. Yang DC, Peters NT, Parzych KR, Uehara T, Markovski M, Bernhardt TG. An ATP-binding cassette transporter-like complex governs cell-wall hydrolysis at the bacterial cytokinetic ring. Proc Natl Acad Sci U S A. 2011;108(45):E1052-60. Epub 20111017. doi: 10.1073/pnas.1107780108. PubMed PMID: 22006326; PubMed Central PMCID: PMCPMC3215046.

48. Du S, Pichoff S, Lutkenhaus J. FtsEX acts on FtsA to regulate divisome assembly and activity. Proc Natl Acad Sci U S A. 2016;113(34):E5052-61. Epub 20160808. doi: 10.1073/pnas.1606656113. PubMed PMID: 27503875; PubMed Central PMCID: PMCPMC5003251.

49. Lee J, Page R, García-Contreras R, Palermino JM, Zhang XS, Doshi O, et al. Structure and function of the Escherichia coli protein YmgB: a protein critical for biofilm formation and acid-resistance. J Mol Biol. 2007;373(1):11–26. Epub 20070802. doi: 10.1016/j.jmb.2007.07.037. PubMed PMID: 17765265; PubMed Central PMCID: PMCPMC2185545.

50. Kettles RA, Tschowri N, Lyons KJ, Sharma P, Hengge R, Webber MA, Grainger DC. The Escherichia coli MarA protein regulates the ycgZ-ymgABC operon to inhibit biofilm formation. Mol Microbiol. 2019;112(5):1609–25. Epub 20190929. doi: 10.1111/mmi.14386. PubMed PMID: 31518447; PubMed Central PMCID: PMCPMC6900184.

51. Palmer AD, Slauch JM. Envelope Stress and Regulation of the Salmonella Pathogenicity Island 1 Type III Secretion System. J Bacteriol. 2020;202(17). Epub 20200810. doi: 10.1128/jb.00272-20. PubMed PMID: 32571967; PubMed Central PMCID: PMCPMC7417839.

52. Shiba Y, Matsumoto K, Hara H. DjlA negatively regulates the Rcs signal transduction system in Escherichia coli. Genes Genet Syst. 2006;81(1):51–6. doi: 10.1266/ggs.81.51. PubMed PMID: 16607041.

53. Jacob-Dubuisson F, Mechaly A, Betton JM, Antoine R. Structural insights into the signalling mechanisms of two-component systems. Nat Rev Microbiol. 2018;16(10):585–93. doi: 10.1038/s41579-018-0055-7. PubMed PMID: 30008469.

54. Mascher T, Helmann JD, Unden G. Stimulus perception in bacterial signal-transducing histidine kinases. Microbiol Mol Biol Rev. 2006;70(4):910–38. doi: 10.1128/mmbr.00020-06. PubMed PMID: 17158704; PubMed Central PMCID: PMCPMC1698512.

55. Krell T, Lacal J, Busch A, Silva-Jiménez H, Guazzaroni ME, Ramos JL. Bacterial sensor kinases: diversity in the recognition of environmental signals. Annu Rev Microbiol. 2010;64:539–59. doi: 10.1146/annurev.micro.112408.134054. PubMed PMID: 20825354.

56. Schumacher K, Gelhausen R, Kion-Crosby W, Barquist L, Backofen R, Jung K. Ribosome profiling reveals the fine-tuned response of Escherichia coli to mild and severe acid stress. mSystems. 2023;8(6):e0103723. Epub 20231101. doi: 10.1128/msystems.01037-23. PubMed PMID: 37909716; PubMed Central PMCID: PMCPMC10746267.

57. Flores-Kim J, Darwin AJ. The Phage Shock Protein Response. Annu Rev Microbiol. 2016;70:83–101. Epub 20160608. doi: 10.1146/annurev-micro-102215-095359. PubMed PMID: 27297125.

58. Todor H, Herrera N, Gross CA. Three Bacterial DedA Subfamilies with Distinct Functions and Phylogenetic Distribution. mBio. 2023;14(2):e0002823. Epub 20230301. doi: 10.1128/mbio.00028-23. PubMed PMID: 36856409; PubMed Central PMCID: PMCPMC10127716.

59. Baba T, Ara T, Hasegawa M, Takai Y, Okumura Y, Baba M, et al. Construction of Escherichia coli K-12 in-frame, single-gene knockout mutants: the Keio collection. Mol Syst Biol. 2006;2:2006.0008. Epub 20060221. doi: 10.1038/msb4100050. PubMed PMID: 16738554; PubMed Central PMCID: PMCPMC1681482.

60. Gibson DG, Young L, Chuang RY, Venter JC, Hutchison CA, 3rd, Smith HO. Enzymatic assembly of DNA molecules up to several hundred kilobases. Nat Methods. 2009;6(5):343–5. Epub 20090412. doi: 10.1038/nmeth.1318. PubMed PMID: 19363495.

61. Ulbrandt ND, Newitt JA, Bernstein HD. The E. coli signal recognition particle is required for the insertion of a subset of inner membrane proteins. Cell. 1997;88(2):187–96. doi: 10.1016/s0092-8674(00)81839-5. PubMed PMID: 9008159.

