## Supplementary material for "RcsF-independent mechanisms of signaling within the Rcs Phosphorelay": Petchiappan et al Supplemental Figs. and Tables

**Supplementary Figures, S1-S9**

**Supplementary Tables:**

**S1: Strains**

**S2: Plasmids**

**S3: Primers**

**S4: gBlocks**

**References**

**Supplementary Figures:**

**Fig. S1:**

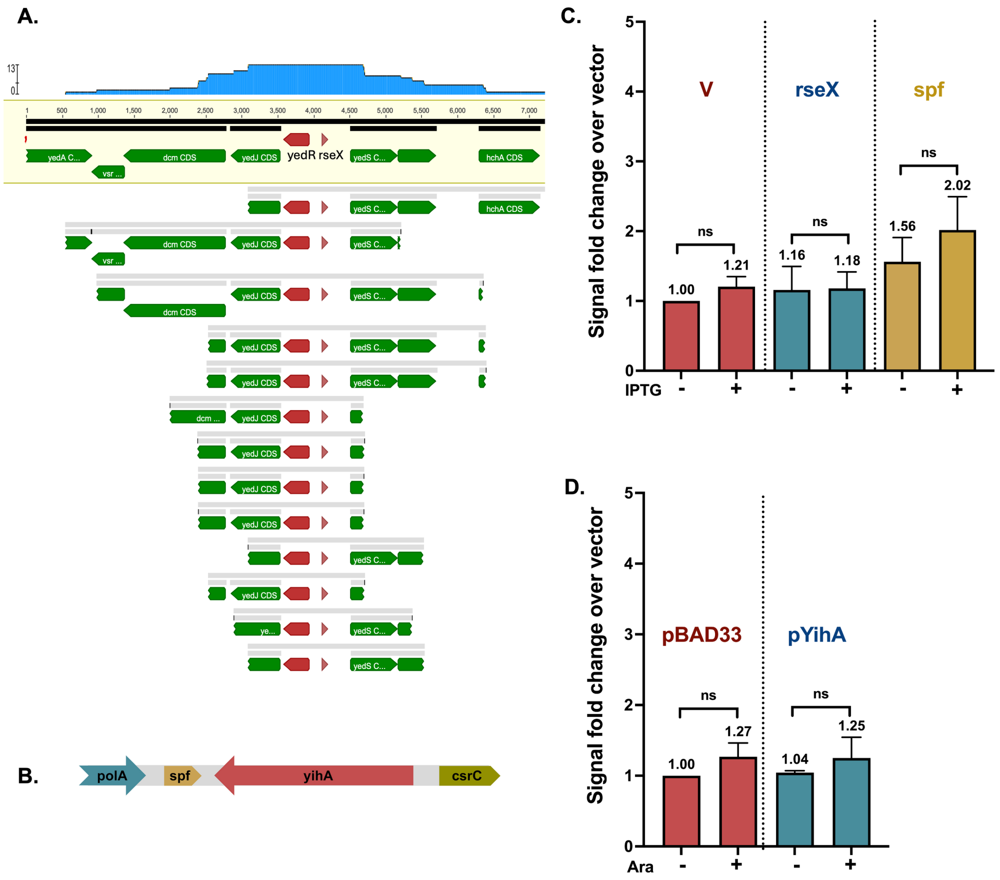

**Fig. S2:**

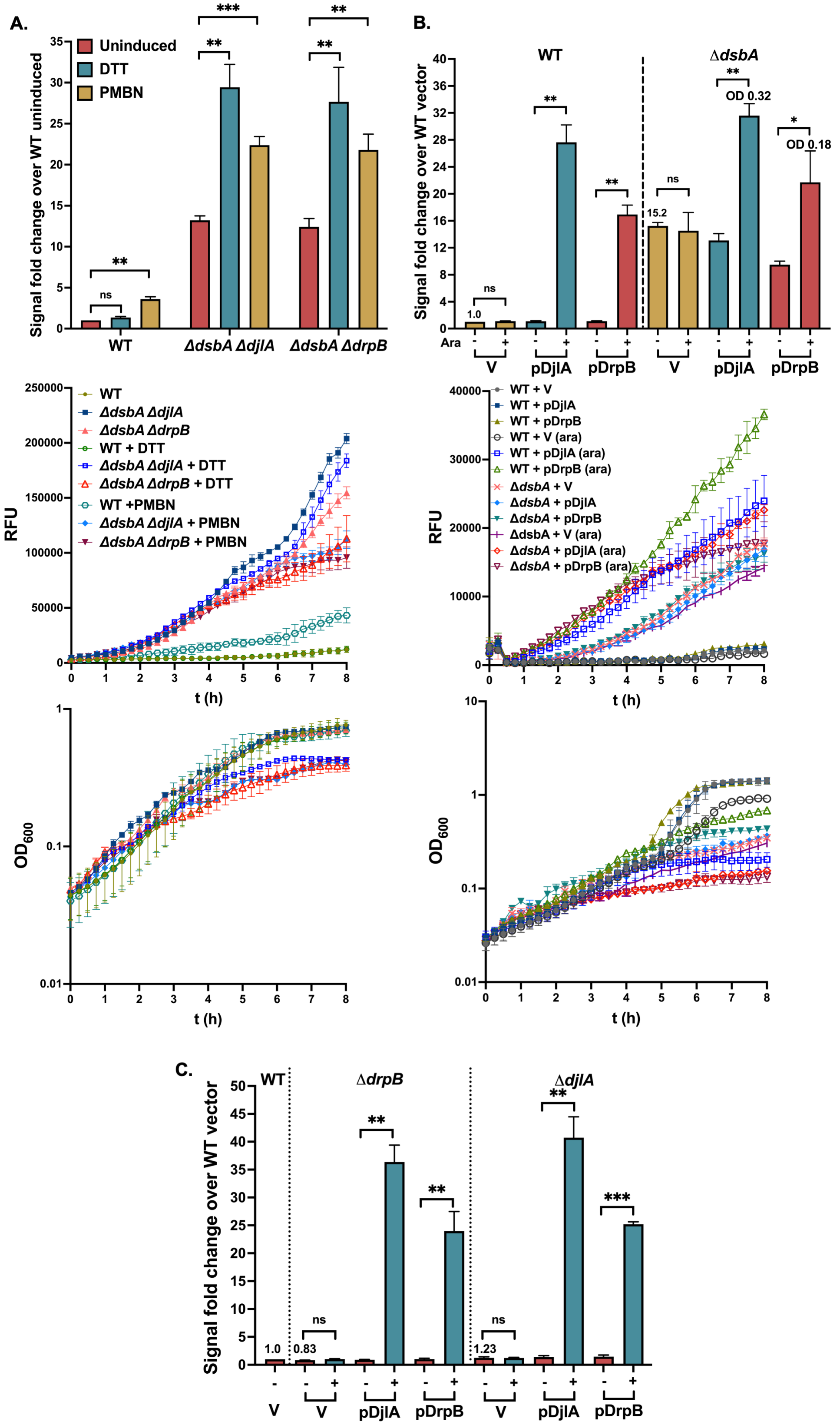

A.
Uninduced
DTT
PMBN
Signal fold change over WT uninduced
WT
ΔdsbA ΔdjlA
ΔdsbA ΔdrpB
ns
**
***
B.
WT
ΔdsbA
Signal fold change over WT vector
Ara
V
pDjlA
pDrpB
1.0
15.2
OD 0.32
OD 0.18
RFU
t (h)
WT
ΔdsbA ΔdjlA
ΔdsbA ΔdrpB
WT + DTT
ΔdsbA ΔdjlA + DTT
ΔdsbA ΔdrpB + DTT
WT +PMBN
ΔdsbA ΔdjlA + PMBN
ΔdsbA ΔdrpB + PMBN
WT + V
WT + pDjlA
WT + pDrpB
WT + V (ara)
WT + pDjlA (ara)
WT + pDrpB (ara)
ΔdsbA + V
ΔdsbA + pDjlA
ΔdsbA + pDrpB
ΔdsbA + V (ara)
ΔdsbA + pDjlA (ara)
ΔdsbA + pDrpB (ara)
OD600
C.
WT
ΔdrpB
ΔdjlA
1.0
0.83
1.23

**Fig. S3:**

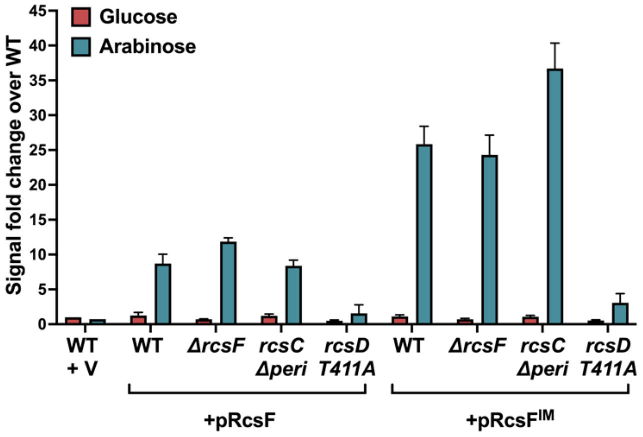
Glucose
Arabinose
Signal fold change over WT
45
40
35
30
25
20
15
10
5
0
WT
+ V
WT
ΔrcsF
rcsC
Δperi
rcsD
T411A
WT
ΔrcsF
rcsC
Δperi
rcsD
T411A
+pRcsF
+pRcsF^IM

**Fig. S4:**

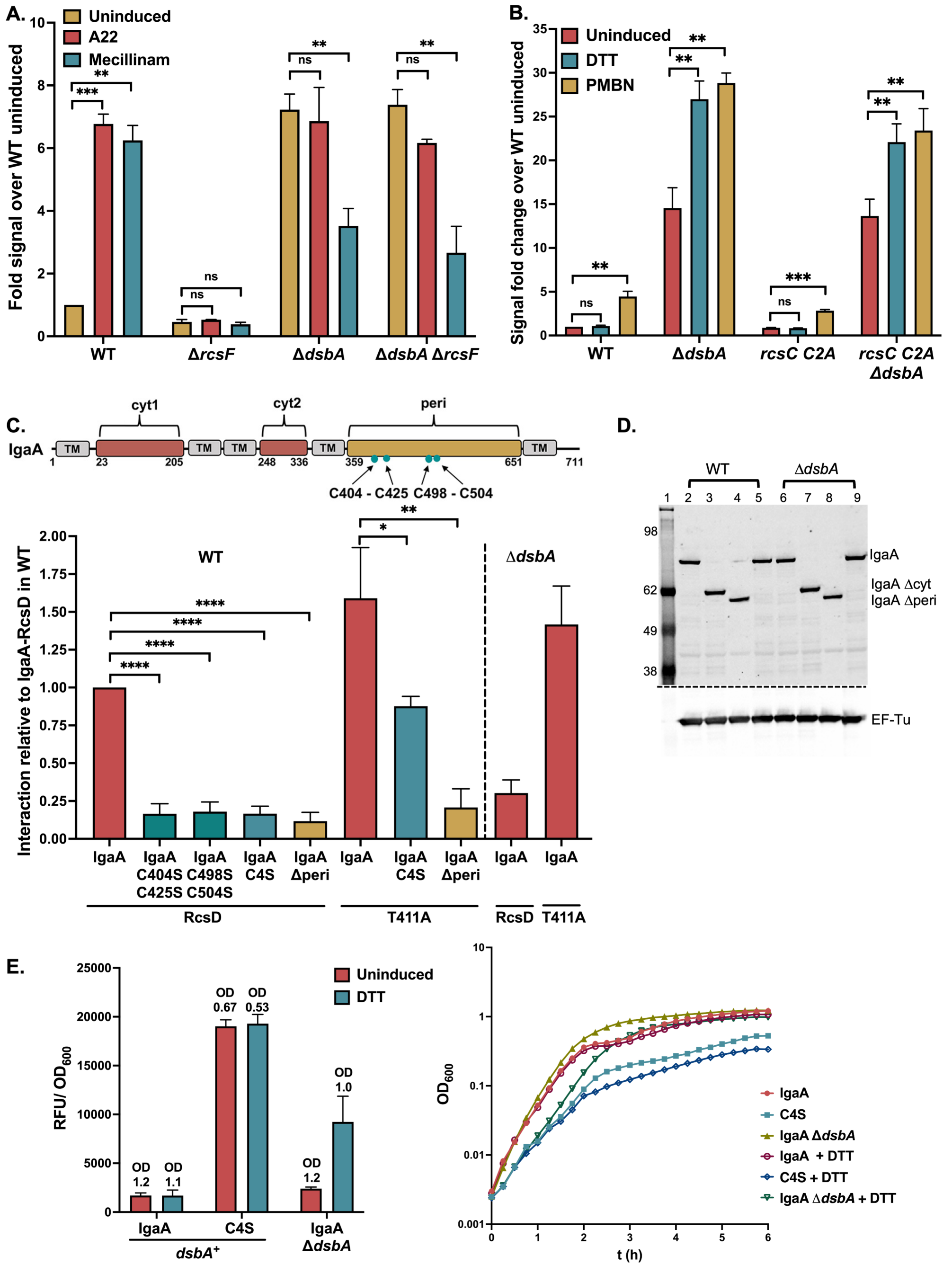
A.
Uninduced
A22
Mecillinam
Fold signal over WT uninduced
***
**
ns
WT
ΔrcsF
ΔdsbA
ΔdsbA ΔrcsF
B.
Uninduced
DTT
PMBN
Signal fold change over WT uninduced
WT
ΔdsbA
rcsC C2A
rcsC C2A ΔdsbA
C.
IgaA
cyt1
cyt2
peri
TM
C404 - C425 C498 - C504
Interaction relative to IgaA-RcsD in WT
WT
ΔdsbA
IgaA
IgaA C404S C425S
IgaA C498S C504S
IgaA C4S
IgaA Δperi
RcsD
T411A
D.
WT
ΔdsbA
IgaA
IgaA Δcyt
IgaA Δperi
EF-Tu
E.
Uninduced
DTT
RFU/ OD600
OD 1.2
OD 1.1
OD 0.67
OD 0.53
OD 1.2
OD 1.0
IgaA
C4S
dsbA+
IgaA ΔdsbA
OD600
t (h)
IgaA
C4S
IgaA ΔdsbA
IgaA + DTT
C4S + DTT
IgaA ΔdsbA + DTT

**Fig. S5:**

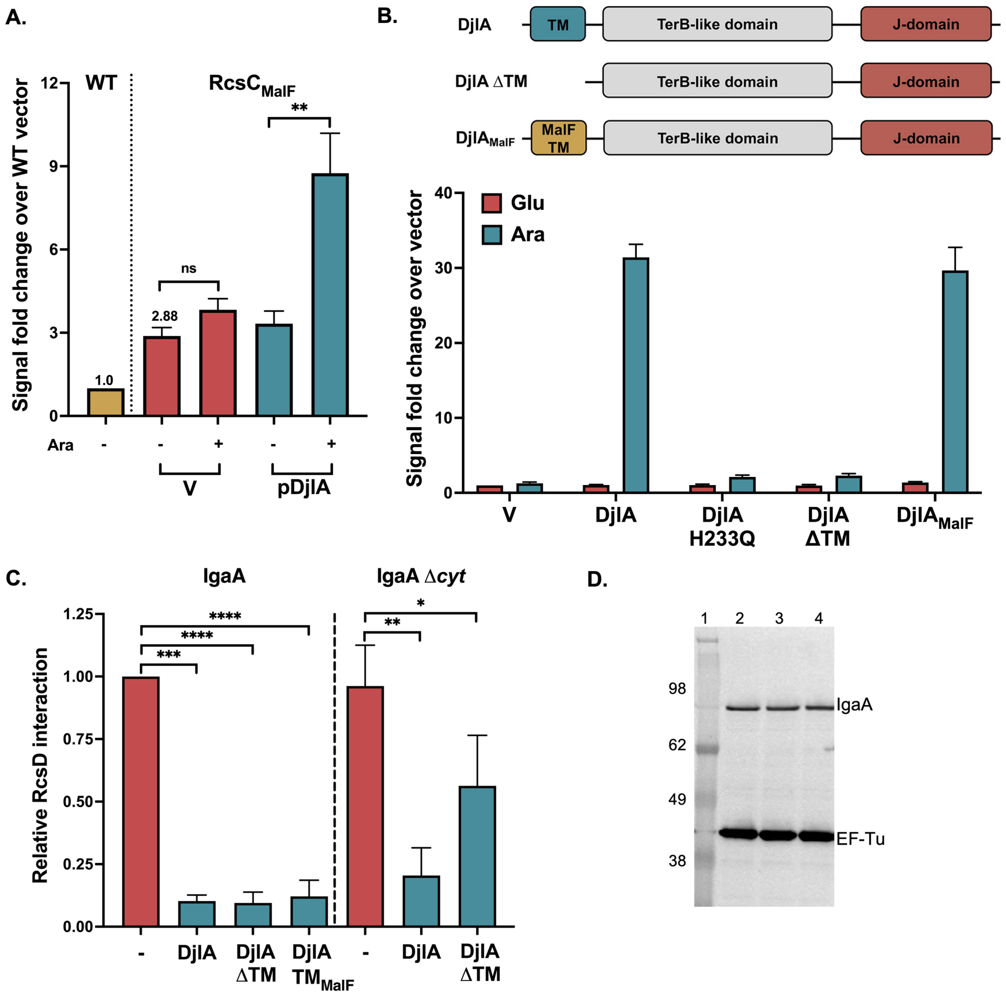
A.
WT
RcsC$_{MalF}$
Signal fold change over WT vector
12
9
6
3
0
1.0
2.88
ns
**
Ara
-
-
+
-
+
V
pDjlA
B.
DjlA
TM
TerB-like domain
J-domain
DjlA ΔTM
TerB-like domain
J-domain
DjlA$_{MalF}$
MalF TM
TerB-like domain
J-domain
Glu
Ara
Signal fold change over vector
40
30
20
10
0
V
DjlA
DjlA H233Q
DjlA ΔTM
DjlA$_{MalF}$
C.
IgaA
IgaA Δ*cyt*
Relative RcsD interaction
1.25
1.00
0.75
0.50
0.25
0.00
****
****
***
*
**
-
DjlA
DjlA ΔTM
DjlA TM$_{MalF}$
-
DjlA
DjlA ΔTM
D.
1
2
3
4
98
62
49
38
IgaA
EF-Tu

**Fig. S6:**

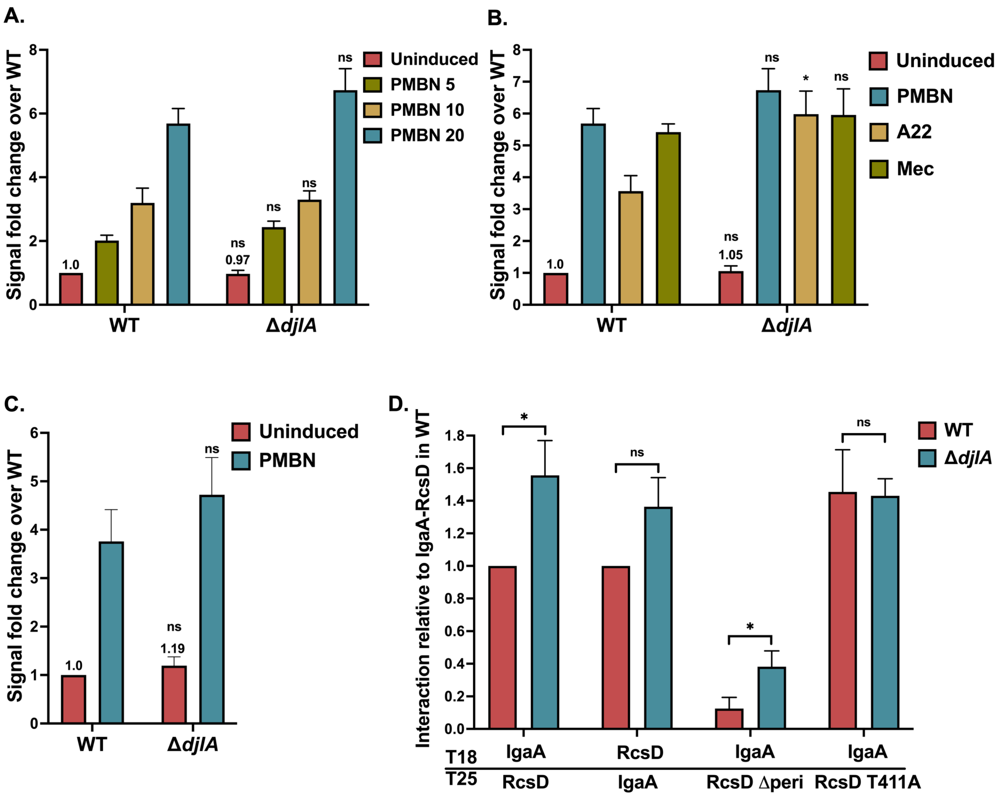
A.
Signal fold change over WT
Uninduced
PMBN 5
PMBN 10
PMBN 20
1.0
0.97
ns
WT
ΔdjlA
B.
Signal fold change over WT
Uninduced
PMBN
A22
Mec
1.0
1.05
ns
*
WT
ΔdjlA
C.
Signal fold change over WT
Uninduced
PMBN
1.0
1.19
ns
WT
ΔdjlA
D.
Interaction relative to IgaA-RcsD in WT
WT
ΔdjlA
*
ns
T18
T25
IgaA
RcsD
RcsD
IgaA
IgaA
RcsD Δperi
IgaA
RcsD T411A

**Fig. S7:**

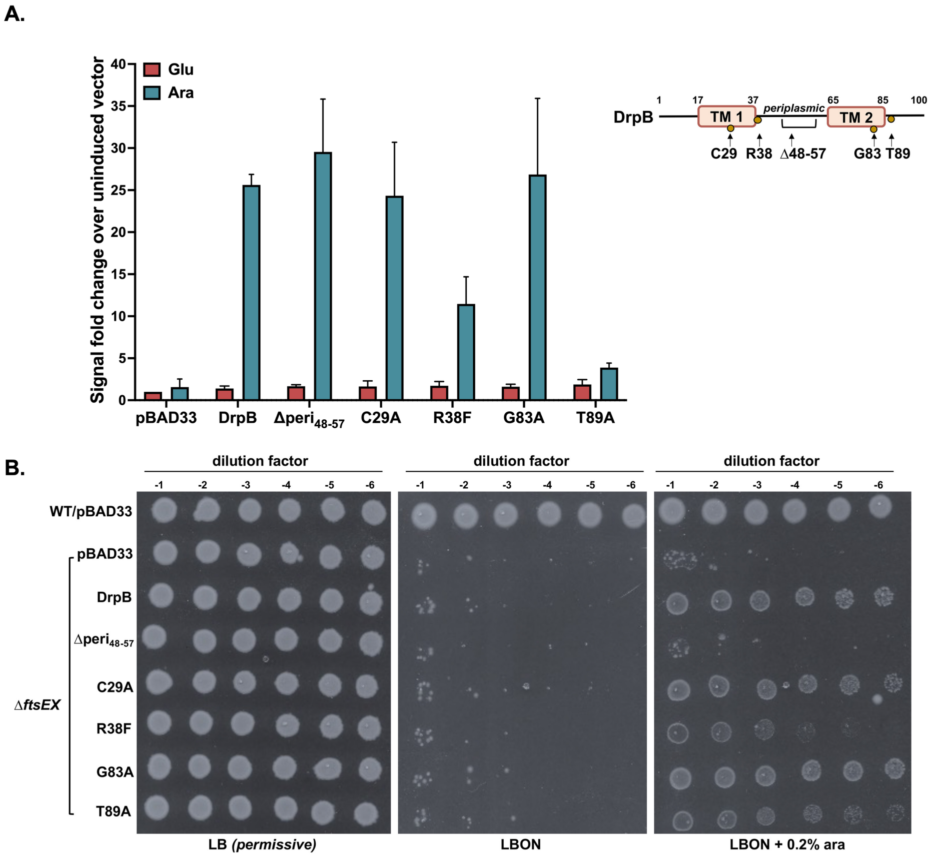
A.
Glu
Ara
Signal fold change over uninduced vector
0
5
10
15
20
25
30
35
40
pBAD33
DrpB
Δperi48-57
C29A
R38F
G83A
T89A
DrpB
1
17
37
65
85
100
TM 1
periplasmic
TM 2
C29
R38
Δ48-57
G83
T89
B.
dilution factor
-1
-2
-3
-4
-5
-6
WT/pBAD33
pBAD33
DrpB
Δperi48-57
C29A
R38F
G83A
T89A
ΔftsEX
LB (permissive)
LBON
LBON + 0.2% ara

**Fig. S8:**

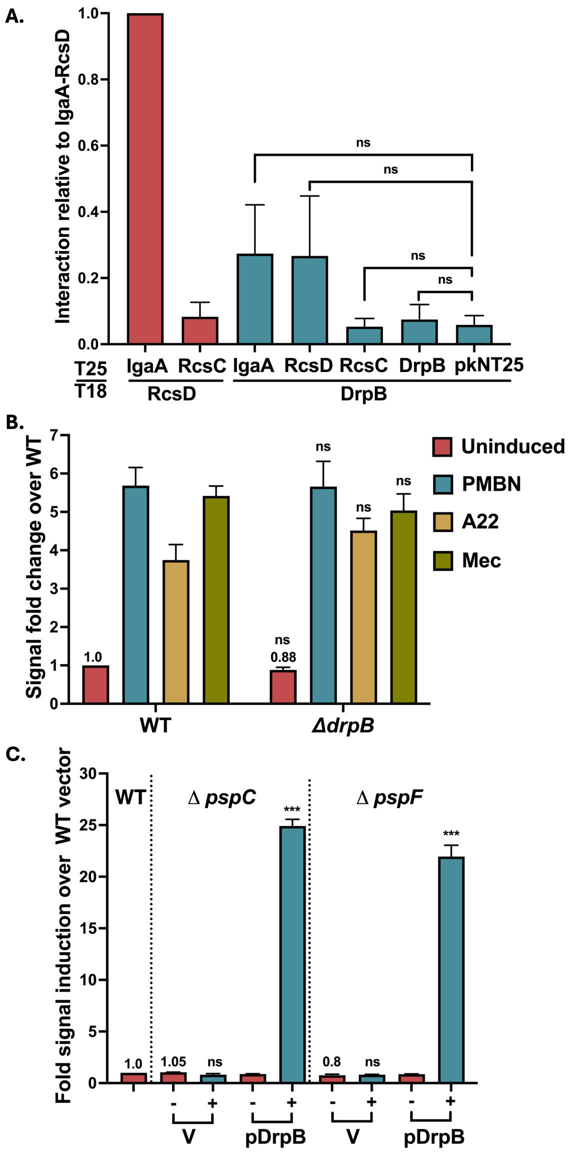

A.
Interaction relative to IgaA-RcsD
1.0
0.8
0.6
0.4
0.2
0.0
ns
ns
ns
ns
T25
T18
IgaA
RcsC
IgaA
RcsD
RcsC
DrpB
pkNT25
RcsD
DrpB
B.
Signal fold change over WT
7
6
5
4
3
2
1
0
1.0
ns
0.88
ns
ns
ns
Uninduced
PMBN
A22
Mec
WT
ΔdrpB
C.
Fold signal induction over WT vector
30
25
20
15
10
5
0
WT
Δ pspC
Δ pspF
***
***
1.0
1.05
ns
0.8
ns
-
+
V
pDrpB
V
pDrpB

**Fig. S9:**

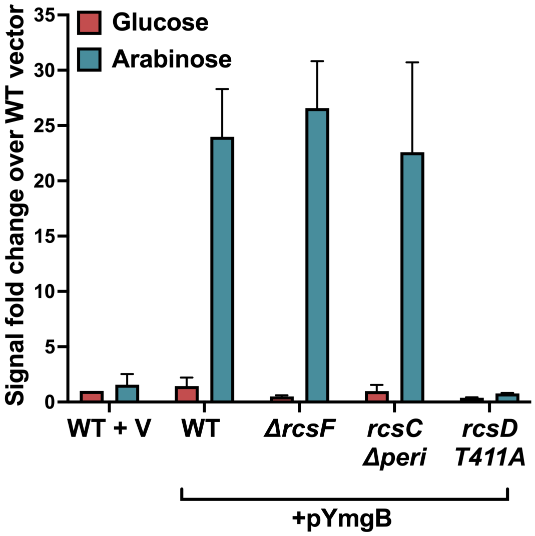
Glucose
Arabinose
Signal fold change over WT vector
35
30
25
20
15
10
5
0
WT + V
WT
ΔrcsF
rcsC Δperi
rcsD T411A
+pYmgB

<u>**Supplementary Tables**</u>

**Table S1: List of strains used in this study**

Strains were constructed by recombineering or P1 transduction with selectable markers (Table S1). Recombineering was done in strains carrying a chromosomal mini-λ Red system (*miniλ::tet*) or a plasmid-borne Red system (pSIM27). Some strains were generated by direct P1 transduction from the corresponding mutant strains in the Keio collection [1].

| Name | Genotype | Method of construction or reference |
|---|---|---|
| MG1655 | Wild-type *E. coli* K-12 | Lab collection |
| BTH101 | *F-, cya-99, araD139, galE15, galK16, rpsL1 (Str$^r$), hsdR2, mcrA1, mcrB1* | [2] |
| NEB DH5-alpha F'IQ | F´ *proA$^+$B$^+$ lacI$^q$ Δ(lacZ)M15 zzf::Tn10* (Tet$^R$) / *fhuA2Δ(argF-lacZ)U169 phoA glnV44 Φ80Δ(lacZ)M15 gyrA96 recA1 relA1 endA1 thi-1 hsdR17* | New England Biolabs |
| SG20382 | *rcsB11*::Tn10 (*tet$^r$*) | [3] |
| DH300 | P$_{rprA142}$-*lacZ* | [4] |
| DH311 | P$_{rprA142}$-*lacZ*, *rcsB311*::*kan* | [4] |
| DH339 | P$_{rprA142}$-*lacZ*, *yojN*::*kan* (*rcsD542*) | [5] |
| DH375 | P$_{rprA142}$-*lacZ*, *rcsC C111A C154A with atoS*::*kan* | DH300 + P1 (NM344a)$^{\#}$ |
| DJ480 | MG1655 *lacX74* | [6] |
| TKC | *tetA, cat, kan* | [7] |
| NC397 | *W3110 pglΔ8 gal$_{490}$ λ cI857(cro-bioA)$_{Δ}$ lacI$_{◦}$ <>kan-Ter<>cat sacB <>lacZYA* | [8] |
| HK307 | MC1000 *dsbA::kan* | Beckwith lab |
| EC251 | MG1655 | [9] |
| EC855 | MG1655 *ftsE::kan* | Weiss lab |
| EC1215 | *ΔftsEX<>frt* | [9] |
| NM7 | *ΔrcsF12*::*cat-sacB* | [10] |
| NM300 | DJ480 *mini-λ-tet* | [11] |
| NM338 | DJ480 *rcsC C111A- cat-sacB* | NM300 + linear transformation |

| | | |
|---|---|---|
| NM340 | DJ480 *rcsC C111A* | NM338 + single-stranded Cys111Ala replacement primer |
| NM344a | DJ480 *rcsC C111A C154A with atoS*::*kan* | NM355 + linear transformation |
| NM350 | DJ480 *rcsC C111A with atoS*::*kan* | NM340 + linear transformation |
| NM355 | DJ480 *ΔrcsC154-atoS::cat* | NM300 linear transformation |
| NM358 | DJ480 *mini-λ-tet, rcsB311::kan* | NM300 + P1 (DH311) |
| NM364 | DJ480 *mini-λ-tet, rcsB311::kan, Δwza-ΔcpsB::zeo* | NM358 electroporated with NM1201 PCR product (cpsB-zeo.R and wza-zeo.F) |
| NM1201 | MG1655 *ybeW::zeo* | [12] |
| EAW1 | *rcsB11*::Tn10, *cya* | [10] |
| EAW2 | *rcsC32*::Tn10, *cya* | [10] |
| EAW4 | *ΔrcsF12*::*cat-sacB*, *cya* | [10] |
| EAW8 | *ΔaraBAD*::$P_{rprA142}$-mCherry, *ΔaraE*p $P_{CP6}$::*gent*::$P_{cp18}$-*araE* | [10] |
| EAW18 | *ΔaraBAD*::$P_{rprA142}$-mCherry, *rcsC*::Tn10, *ΔaraE*p $P_{cp6}$*gent*::$P_{cp18}$-*araE* | [10] |
| EAW19 | *ΔaraBAD*::$P_{rprA142}$-mCherry, *rcsD*541(::FRT), *ΔaraEp* $P_{cp6}$*gent*::$P_{cp18}$-*araE* | [10] |
| EAW25 | *ΔaraBAD*::$P_{rprA142}$-mCherry, *ΔaraE*p $P_{CP6}$::*gent*::$P_{cp18}$-*araE, Δwza-ΔcpsB::zeo* | EAW8 + P1 from NM364 (screened for *Δwza-ΔcpsB::zeo*) |
| EAW31 | *ΔaraBAD*::$P_{rprA142}$-mCherry, *ΔaraE*p $P_{cp6}$*gent*::$P_{cp18}$-*araE, rcsB*::*kan* | [10] |
| EAW32 | *ΔaraBAD*::$P_{rprA142}$-mCherry, *ΔaraE*p $P_{cp6}$*gent*::$P_{cp18}$-*araE*, D*rcsF12*::*cat-sacB* | [10] |
| EAW34 | *ΔaraBAD*::$P_{rprA142}$-mCherry, *ΔaraE*p $P_{CP6}$::*gent*::$P_{cp18}$-*araE, Δwza-ΔcpsB::zeo*, *ΔrcsF12::cat-sacB* | EAW25 + P1 (NM7) |
| EAW70 | *ΔaraBAD*::$P_{rprA142}$-mCherry, *ΔaraE*p $P_{cp6}$*gent*::$P_{cp18}$-*araE*, *ΔrcsC51::rcsCΔ48-314* | [10] |
| EAW72 | *ΔaraBAD*::$P_{rprA142}$-mCherry, *ΔaraE*p $P_{cp6}$*gent*::$P_{cp18}$-*araE*, *ΔrcsC51::*$rcsC_{1-19}$-$malF_{2-59}$-$rcsC_{334-C}$ | [10] |
| EAW120 | *ΔaraBAD*::$P_{rprA142}$-mCherry, *ΔaraE*p $P_{cp6}$*gent*::$P_{cp18}$-*araE*, *rcsD841** | [10] |

| | | |
|---|---|---|
| EAW121 | *ΔaraBAD*::$P_{rprA142}$-mCherry, *ΔaraE*p $P_{cp6}$*gent*::$P_{cp18}$-*araE*, *rcsDT411A* | [10] |
| EAW62 | *ΔaraBAD*::$P_{rprA142}$-mCherry, *ΔaraE*p $P_{CP6}$::*gent*::$P_{cp18}$-*araE*, *dsbA::kan* | EAW8 + P1 (HK307) |
| EAW63 | *ΔaraBAD*::$P_{rprA142}$-mCherry, *rcsC*::Tn10, *ΔaraE*p $P_{cp6}$*gent*::$P_{cp18}$-*araE*, *dsbA::kan* | EAW18 + P1 (HK307) |
| EAW67 | *ΔaraBAD*::$P_{rprA142}$-mCherry, *ΔaraE*p $P_{CP6}$::*gent*::$P_{cp18}$-*araE*, *dsbA::kan*, Δ*rcsF12::cat-sacB* | EAW62 + P1 (NM7) |
| EAW74 | *ΔaraBAD*::$P_{rprA142}$-mCherry, *ΔaraE*p $P_{cp6}$*gent*::$P_{cp18}$-*araE*, *ΔrcsC51::rcsCΔ48-314*, *dsbA::kan* | EAW70 + P1 (HK307) |
| EAW90 | *ΔaraBAD*::$P_{rprA142}$-mCherry, *ΔaraE*p $P_{cp6}$*gent*::$P_{cp18}$-*araE*, *rcsD541(::FRT)*, *ΔigaA*::*kan-araC-kid* | [10] |
| AP11 | *ΔaraBAD*::$P_{rprA142}$-mCherry, *ΔaraE*p $P_{CP6}$::*gent*::$P_{cp18}$-*araE*, Δ*dsbA* | EAW62 + pcp20 |
| AP12 | *ΔaraBAD*::$P_{rprA142}$-mCherry, *ΔaraE*p $P_{CP6}$::*gent*::$P_{cp18}$-*araE*, Δ*dsbA*, *rcsB311::kan* | AP11 + P1 (DH311) |
| AP13 | *ΔaraBAD*::$P_{rprA142}$-mCherry, *ΔaraE*p $P_{CP6}$::*gent*::$P_{cp18}$-*araE*, Δ*dsbA*, *yojN::kan (rcsD542)* | AP11 + P1 (DH339) |
| AP14 | *ΔaraBAD*::$P_{rprA142}$-mCherry, *ΔaraE*p $P_{cp6}$*gent*::$P_{cp18}$-*araE*, *rcsDT411A*, *dsbA::kan* | EAW121 + P1 (HK307) |
| AP41 | *ΔaraBAD*::$P_{rprA142}$-mCherry, *ΔaraE*p $P_{CP6}$::*gent*::$P_{cp18}$-*araE*, *drpB::kan* | EAW8 + P1 (Keio JW1946) |
| AP46 | *ΔaraBAD*::$P_{rprA142}$-mCherry, *ΔaraE*p $P_{CP6}$::*gent*::$P_{cp18}$-*araE*, *djlA::kan* | EAW8 + P1 (Keio JW0054) |
| AP50 | *ΔaraBAD*::$P_{rprA142}$-mCherry, *ΔaraE*p $P_{CP6}$::*gent*::$P_{cp18}$-*araE*, *rcsB11::Tn10 (tetr)* | EAW8 + P1 (SG20382) |
| AP51 | D*araBAD*::$P_{rprA142}$-mCherry, D*araE*p $P_{CP6}$::*gent*::$P_{cp18}$-*araE*, *rcsF::kan* | EAW8 + P1 (Keio JW0192) |
| AP57 | *dsbA::kan, cya* | BTH101 + P1 (HK307) |
| AP58 | *ΔdsbA, cya* | AP57 + pcp20 |
| AP71 | *ΔaraBAD*::$P_{rprA142}$-mCherry, *ΔaraE*p $P_{CP6}$::*gent*::$P_{cp18}$-*araE*, Δ*dsbA*, *drpB::kan* | AP11 + P1 (Keio JW1946) |
| AP72 | *ΔaraBAD*::$P_{rprA142}$-mCherry, *ΔaraE*p $P_{CP6}$::*gent*::$P_{cp18}$-*araE*, Δ*dsbA*, *djlA::kan* | AP11 + P1 (Keio JW0054) |
| AP113 | *ΔaraBAD*::$P_{rprA142}$-mCherry, *ΔaraE*p $P_{CP6}$::*gent*::$P_{cp18}$-*araE*, *pspC::kan* | EAW8 + P1 (Keio JW1299) |

| | | |
|---|---|---|
| AP114 | *ΔaraBAD*::$P_{rprA142}$-mCherry, *ΔaraE*p $P_{CP6}$::*gent*::$P_{cp18}$-*araE, pspF::kan* | EAW8 + P1 (Keio JW1296) |
| AP154 | *ΔaraBAD*::$P_{rprA142}$-mCherry, *ΔaraE*p $P_{CP6}$::*gent*::$P_{cp18}$-*araE, ftsE::kan* | EAW8 + P1 (EC855) |
| AP155 | *ΔaraBAD*::$P_{rprA142}$-mCherry, *ΔaraE*p $P_{CP6}$::*gent*::$P_{cp18}$-*araE, rcsB11::Tn10 (tet$^r$), ftsE::kan* | AP50 + P1 (EC855) |
| AP158 | *rcsB311*::*kan* | EC251 + P1 (DH311) |
| AP159 | *ΔftsEX<>frt, rcsB311*::*kan* | EC1215 + P1 (DH311) |
| AP168 | *ΔaraBAD*::$P_{rprA142}$-mCherry, *ΔaraE*p $P_{cp6}$*gent*::$P_{cp18}$-*araE*, *rcsD541(::FRT)*, *ΔigaA::igaA C404S C424S C498S C504S* | EAW90 recombination with PCR product of pEAW1C4S template using oligos EAW213 and EAW214 |
| AP169 | *ΔaraBAD*::$P_{rprA142}$-mCherry, *rcsD541(::FRT)*, *ΔaraEp* $P_{cp6}$*gent::*$P_{cp18}$-*araE*, *dsbA::kan* | EAW19 + P1 (EAW62) |
| AP172 | *ΔaraBAD*::$P_{rprA142}$-mCherry, *ΔaraE*p $P_{CP6}$::*gent*::$P_{cp18}$-*araE, rcsC C111A C154A with atoS::kan* | EAW8 + P1 (DH375) |
| AP173 | *ΔaraBAD*::$P_{rprA142}$-mCherry, *ΔaraE*p $P_{CP6}$::*gent*::$P_{cp18}$-*araE, rcsC C111A C154A with atoS::kan, ΔdsbA* | AP11 + P1 (DH375) |

**[#]Construction of NM344a:**

NM344a was constructed in two steps. In the first step, the region between *atoS* and *rcsCcys154* in NM300 was replaced by recombineering a *cat* resistance cassette using the primers Δ*atoS-rcsC154.CmF* and Δ*atoS-rcsC154.CmR* and the TKC strain used as a template. This generated NM355. In a second parallel step, NM338 was generated by recombineering the PCR product (primers Cys111-ala_cat and Cys111-ala_sacB) from NC397 into NM300. Next, NM340 was generated by linear transformation of the single-stranded oligo Cys111Ala replacement primer into NM338. In this strain NM340, a *kan* resistance cassette was then inserted between *rcsCcys111* and *atoS* using primers RcsC-KAN-AtoS.F and RcsC-KAN-AtoS.R. This generated strain NM350. A 2-kb fragment from the *kan* cassette to the rcsC154 nucleotide was amplified from NM350 using the primers atoS_RcsCys154 (containing the Cys154Ala mutation) and RcsC-KAN-AtoS.R was used to transform NM355. This generated strain NM344a.

**Table S2: List of plasmids used in this study**

Plasmids were constructed by the Gibson assembly method using the In-fusion HD Cloning kit (Takara Bio USA) [13]. Site-directed mutagenesis (SDM) in the genes was carried out using the QuikChange Site-directed mutagenesis kit (Agilent).

| Name | Description | Method of construction/ Reference |
|---|---|---|
| pBAD24 | Vector for protein expression regulated by the arabinose operon (Amp$^r$) | [14] |
| pBAD33 | Vector for protein expression regulated by the arabinose operon (Chl$^r$) | [14] |
| pCP20 | Plasmid with temperature-sensitive origin of replication, encoding the FLP recombinase | [15] |
| pUT18 | Vector encoding the Cya T18 fragment under lac promoter control (Amp$^r$) | [16] |
| pKNT25 | Vector encoding the Cya T18 fragment under lac promoter control (Kan$^r$) | [17] |
| pSIM27 | Plasmid with temperature-sensitive origin of replication, encoding l-Red cI857, gam-beta-exo | https://ncifrederick.cancer.gov/recombineering/strains-plasmids-and-primers |
| pEAW1 | IgaA with T18 tag at C-terminal cloned in pUT18 | [10] |
| pEAW1cyt1 | IgaA *Δ*36-181 with T18 tag at C-terminal cloned in pUT18 | [10] |
| pEAW1peri | IgaA *Δ*384-649 with T18 tag at C-terminal cloned in pUT18 | [10] |
| pEAW2 | IgaA with T25 tag at C-terminal cloned in pKNT25 | [10] |
| pEAW6 | RcsC with T25 tag at C-terminal cloned in pKNT25 | [10] |
| pEAW7 | RcsD with T18 tag at C-terminal cloned in pUT18 | [10] |
| pEAW8 | RcsD with T25 tag at C-terminal cloned in pKNT25 | [10] |
| pEAW8peri | RcsD *Δ*45-304 with T25 tag at C-terminal cloned in pKNT25 | [10] |
| pEAW8T | RcsD T411A with T25 tag at C-terminal cloned in pKNT25 | [10] |
| pEAW11 | RcsD cloned in pBAD24 | [10] |
| pEAW11T | RcsD T411A cloned in pBAD24 | [10] |

| | | |
|---|---|---|
| pPSG961 | DjlA cloned in pBAD33 | [18] (Gift from A. Jacq?) |
| pDSW1977 | DrpB cloned in pBAD33 | [9] (Gift from David Weiss) |
| pBR-plac | pBR322 derivative plasmid for sRNA overexpression under an artificial $P_{lac}$ promoter | [19] |
| p-rseX | rseX sRNA cloned in pBR-plac | [20] |
| p-spf | spot42 sRNA cloned in pBR-plac | [20] |
| pNM654 | IgaA cloned in pBAD24 | IgaA amplified from genomic DNA (yrfF_pBAD24F and R); pBAD24 digested with *EcoRI* and *HindIII* |
| pNM656 | IgaA C425S cloned in pBAD24 | pNM654 template with primers yrfF Cys425Ser.F and Cys425Ser.R (SDM) |
| pNM665 | IgaA C425S C498S C504S cloned in pBAD24 | pNM656 template with primers yrfF Cys498-504Ser.F and Cys498-504Ser.R (SDM) |
| pNM671 | IgaA C404S C425S C498S C504S cloned in pBAD24 | pNM665 template with primers yrfF Cys404Ser.F and Cys404Ser.R (SDM) |
| pAP101 | IgaA *Δ*36-181 Δ263-329 with T18 tag at C-terminal cloned in pUT18 | pEAW1cyt1 template with primers EW209 and EW210 |
| pAP102 | IgaA C404S C425S with T18 tag at C-terminal cloned in pUT18 | pAP104 template with primers AP691 and AP692 (SDM) |
| pAP103 | IgaA C498S C504S with T18 tag at C-terminal cloned in pUT18 | pAP105 template with primers AP587 and AP588 (SDM) |
| pAP104 | IgaA C404S with T18 tag at C-terminal cloned in pUT18 | pEAW1 template with primers AP351 and AP352 (SDM) |
| pAP105 | IgaA C498S with T18 tag at C-terminal cloned in pUT18 | pEAW1 template with primers AP693 and AP694 (SDM) |
| pEAW1C4S | IgaA C404S C425S C498S C504S (C4S) with T18 tag at C-terminal cloned in pUT18 | Insert from pNM671 (EW6 and EW7); pUT18 linearized with EW1 and EW2 |
| pAP1401 | IgaA with T18 tag at C-terminal and DjlA cloned downstream under the same promoter in pUT18 | Insert from gBlock AP_GJ1; pEAW1 linearized with AP375 and AP376 |
| pAP1402 | IgaA with T18 tag at C-terminal and DjlA H233Q cloned downstream under the same promoter in pUT18 | pAP1401 template with primers AP459 and AP460 (SDM) |
| pAP1403 | IgaA with T18 tag at C-terminal and DjlA Δ1-31 (ΔTM) cloned downstream under the same promoter in pUT18 | pAP1401 template with primers AP495 and AP496 (SDM) |

| | | |
|---|---|---|
| pAP1404 | IgaA with T18 tag at C-terminal and DjlA $TM_{MalF}$ cloned downstream under the same promoter in pUT18 | Insert from gBlock AP_GJMF; pEAW1 linearized with AP375 and AP376 |
| pAP407 | DrpB with T18 tag at C-terminal cloned in pUT18 | Insert from pDSW1977 (AP241 and AP242); pUT18 linearized with EW1 and EW2 |
| pAP408 | DrpB with T25 tag at C-terminal cloned in pKNT25 | Insert from pDSW1977 (AP241 and AP242); pUT18 linearized with EW1 and EW2 |
| pAP804 | RcsD T411A Δ45-304 with T25 tag at C-terminal cloned in pKNT25 | pEAW8peri template with primers T411A F and T411A R (SDM) |
| pAP3301 | DrpB Δ48-59 cloned in pBAD33 | pDSW1977 template with primers AP321 and AP322 (SDM) |
| pAP3304 | DrpB C29A cloned in pBAD33 | pDSW1977 template with primers AP323 and AP324 (SDM) |
| pAP3305 | DrpB R38F cloned in pBAD33 | pDSW1977 template with primers AP325 and AP326 (SDM) |
| pAP3306 | DrpB G83A cloned in pBAD33 | pDSW1977 template with primers AP327 and AP328 (SDM) |
| pAP3307 | DrpB T89A cloned in pBAD33 | pDSW1977 template with primers AP329 and AP330 (SDM) |
| pAP3311 | DjlA Δ1-31 (ΔTM) cloned in pBAD33 | pPSG961 template with primers AP563 and AP564 |
| pAP3312 | DjlA $TM_{MalF}$ cloned in pBAD33 | Insert from gBlock AP_GJMF (AP565 and AP566); pBAD33 linearized with AP559 and AP560 |
| pAP3315 | DjlA H233Q cloned in pBAD33 | pPSG961 template with primers AP459 and AP460 (SDM) |
| pAP3325 | YmgB/AriR cloned in pBAD33 | Insert from gBlock AP_GymgB; pBAD33 linearized with AP559 and AP560 |
| pAP3327 | YihA cloned in pBAD33 | Insert from gBlock AP_GyihA; pBAD33 linearized with AP559 and AP560 |
| pAP3340 | RcsF cloned in pBAD33 | Insert from gBlock AP_GF33; pBAD33 linearized with AP559 and AP560 |
| pAP3341 | RcsF S17D M18Q cloned in pBAD33 | pAP3340 template with primers AP501 and AP502 (SDM) |

**Table S3: List of primers used in this study**

| Name | Sequence (5'-3') |
|---|---|
| EW1 | AAT CAT GGT CAT AGC TGT TTC CTG TGT GAA ATT G |
| EW2 | AGC TTG CAT GCC TGC AGG TCG AC |
| EW6 | GCT ATG ACC ATG ATT AGC ACC ATT GTG ATT TTT TTA GCT GCT TTG CTG |
| EW7 | GCA GGC ATG CAA GCT TTC GAT AAG GCT TTC TGA AGG GGT GAT C |
| EW209 | TAA CGG AAA GTT TTT ACC GCG CAG ACA ATG AAT TTC CCG C |
| EW210 | AAA AAC TTT CCG TTA CAG CAC TGG CTG CGC |
| EW213 | ACC ACG CCT GAC AGA CTA AGT AAG ATG GGG AAA GCA TGA GCA CCA TTG TGA TTT TTT TAG CTG CTT TGC |
| EW214 | GAC AGG GTA GCA TAA CCT GCC GCG CAA ACG TGT TAT TCG ATA AGG CTT TCT GAA GGG GTG ATC AGT TG |
| *ΔatoS-rcsC154.CmF* | GCGTTCCACTGGCATATCACGCAGACCGAAATTGGCCATAAAATGAGACGTTG ATCGGCACG |
| *ΔatoS-rcsC154.CmR* | GGTTAAGGTGATGATTTCTCGGCGGTGTATCATATTCCAGACCAGCAATAGACAT AAGCGGGC |
| Cys111-ala_cat | CAGGCGGATGTGCCTGCGTTTGAACCGCTGTTTGCCGACTCCGATGCAAAATG AGCAGTTAGTCGGCAC |
| Cys111-ala_sacB | CGCCAATGACTCCAGAGAACCTCGCCAGGTGTTACTCATGTAGACTGCAAAGG GAAAACTGTCCATATG |
| Cys111Ala replacement primer | CAGGCGGATGTGCCTGCGTTTGAACCGCTGTTTGCCGACTCCGATGCTTCCGC AATGAGTAACACCTGGCGAGGTTCTCTGGAGTCATTGGCG |
| RcsC-KAN-AtoS.F | GGTAGCGGTAAAAGCGTGTTACCGCAATGTTCTCTCTTCTGTGTAGGCTGGAGC TGCTTCG |
| RcsC-KAN-AtoS.R | GGTTAAGGTGATGATTTCTCGGCGGTGTATCATATTCCAGATTCCGGGGATCCG TCGACC |
| atoS_RcsCys154 | CGCGTTCCACTGGCATATCACGCAGACCGAAATTGGCCATAGCGAGGTTATCG CTGCCGATTAAAAATAC |
| cpsB-zeo.R | ATT TAC ACC GCG GTT TCG CAT TCA TTG CCT GAT GCG ACG TAA AAA AAG CCC GCT CAT TAG |
| wza-zeo.F | GTG CAC AGG ATA ATT ACT CTG CCA AAG TGA TAA ATA AAC AGT TGA CAA TTA ATC ATC GGC |
| yrfF_pBAD24F | CAG GAG GAA TTC ATG AGC ACC ATT GTG ATT TTT TTA GC |

| | |
|---|---|
| yrfF_pBAD24R | ACA GCC AAG CTT TTA TTC GAT AAG GCT TTC TG |
| yrfF Cys425Ser.F | TTT TTA CCG TTT GAC AGC TCG CAG ATC ATC T |
| yrfF Cys425Ser.R | AGA TGA TCT GCG AGC TGT CAA ACG GTA AAA A |
| yrfF Cys498-504Ser.F | AAG ACA GCG GAT TTA AGT TCT GCC AAA GAT GAC TGA GTG CGA CTG AAA AAT |
| yrfF Cys498-504Ser.R | ATT TTT CAG TCG CAC TCA GTC ATC TTT GGC AGA ACT TAA ATC CGC TGT CTT |
| yrfF Cys404Ser.F | AGC GGT ACG GGA ATG AGT AAT ATT CGA ACT T |
| yrfF Cys404Ser.R | AAG TTC GAA TAT TAC TCA TTC CCG TAC CGC T |
| AP241 | GCT ATG ACC ATG ATT GAA TAC GGT TCG ACA AAG ATG GAA GAG AGA CTC T |
| AP242 | GCA GGC ATG CAA GCT TTC ATA GCG TCT GCT ACG TGC GG |
| AP321 | GGG AAT CCG GCT CAT CAC CCA GAT GTA AC |
| AP322 | ATG AGC CGG ATT CCC TCT ACC GCG GGA AAG TGG TTA GGG |
| AP323 | CAT TGC CCA GAC GAA ATA GCC CGC CCA TGT ATA GAA AGC CCA CAA |
| AP324 | TTG TGG GCT TTC TAT ACA TGG GCG GGC TAT TTC GTC TGG GCA ATG |
| AP325 | CTC ATC ACC CAG ATG TAA AAC GCC ATT GCC CAG ACG AA |
| AP326 | TTC GTC TGG GCA ATG GCG TTT TAC ATC TGG GTG ATG AG |
| AP327 | ACC AGG CAA TGC TGG CTA ACA ATG CCC CG |
| AP328 | CGG GGC ATT GTT AGC CAG CAT TGC CTG GT |
| AP329 | GGA CGA GGT CGG GCG TAC CAG GCA ATG |
| AP330 | CAT TGC CTG GTA CGC CCG ACC TCG TCC |
| AP351 | CGG AAG TTC GAA TAT TAG ACA TTC CCG TAC CGC TA |
| AP352 | TAG CGG TAC GGG AAT GTC TAA TAT TCG AAC TTC CG |
| AP375 | TTA TAT CGA TTG GCG TTC CAC TGC G |
| AP376 | CTA AGT AAT ATG GTG CAC TCT CAG TAC AAT CTG CTC |
| AP459 | CAGCTTATCGGGCTGGTGTTCACTCATCAGCTTACG |
| AP460 | CGTAAGCTGATGAGTGAACACCAGCCCGATAAGCTG |

| | |
|---|---|
| AP495 | CCC TGG ACC AAC GCC TTT CCG TTA CAG CAC TGG CTG CGC AGT AC |
| AP496 | ATG TTT GAT AAA GCC CGT AGC CGT AAA ATG G |
| AP501 | CGA CAG GGG ATC TGC TTA ACT GGT CAC AGC CGC TTA GCA TGA GTG |
| AP502 | CAC TCA TGC TAA GCG GCT GTG ACC AGT TAA GCA GAT CCC CTG TCG |
| AP559 | GAG CTC GAA TTC GCT AGC CCA AAA AAA CG |
| AP560 | AAG CTT GGC TGT TTT GGC GGA TGA G |
| AP563 | GGC TTT ATC AAA CAT ATA TTC CCC AGA TCG ACA CAC GGA TG |
| AP564 | ATG TTT GAT AAA GCC CGT AGC CGT AAA ATG G |
| AP565 | AGC GAA TTC GAG CTC AGC AGG AGG AAT TCA ATG GAT GTC ATT AAA AAG AAA C |
| AP566 | AAA ACA GCC AAG CTT TCA TTT AAA CCC TTT CTG CTG CTT TAT CAG |
| AP587 | CAT TTT TCA GTC GCA CAC TGT CAT CTT TGG CAG AAC A |
| AP588 | TGT TCT GCC AAA GAT GAC AGT GTG CGA CTG AAA AAT G |
| AP691 | CCA GAT GAT CTG CGA GCT GTC AAA CGG TAA AAA AG |
| AP692 | CTT TTT TAC CGT TTG ACA GCT CGC AGA TCA TCT GG |
| AP693 | CAT CTT TGG CAG AAG ATA AAT CCG CTG TCT TCA GTA C |
| AP694 | GTA CTG AAG ACA GCG GAT TTA TCT TCT GCC AAA GAT G |

**TableS4: List of gBlocks used in this study**

| gBlock name | Sequence |
|---|---|
| **AP_GymgB** | AGCGAATTCGAGCTCAGCAGGAGGAATTCAATGCTTGAAGATACTACAATTCAT<br>AATGCAATAACTGATAAAGCGTTAGCAAGTTACTTTCGCAGTTCGGGTAATTTGT<br>TAGAAGAAGAATCAGCAGTGTTAGGGCAGGCTGTCACCAATTTAATGCTTTCAG<br>GCGATAATGTTAATAATAAAAATATTATCTTAAGTCTGATACACTCCTTGGAAAC<br>AACAAGTGATATTCTCAAAGCTGATGTGATTAGAAAAACACTGGAAATCGTGTT<br>GCGATACACAGCTGATGATATGTAAGCTTGGCTGTTTT |
| **AP_GyihA** | AGCGAATTCGAGCTCAGCAGGAGGAATTCAATGACTAATTTGAATTATCAACAGACGCA<br>TTTTGTGATGAGTGCGCCTGATATTCGCCACCTACCTTCCGATACCGGAATTGAAGTG<br>GCTTTTGCAGGCCGTTCCAACGCAGGTAAATCCAGCGCGCTGAACACGCTGACTAA<br>CCAGAAAAGCCTGGCTCGTACCTCAAAAACCCCAGGGCGCACCCAGCTTATCAACC<br>TGTTTGAAGTGGCTGACGGCAAGCGTCTGGTTGACTTGCCTGGGTACGGTTATGCGG<br>AAGTCCCGGAAGAGATGAAGCGCAAATGGCAGCGTGCGCTCGGCGAATACCTCGAA<br>AAACGTCAGAGCCTGCAAGGTCTGGTGGTGCTAATGGATATTCGCCATCCGCTGAAA<br>GATTTGGATCAGCAGATGATTGAGTGGGCGGTAGACAGCAATATCGCCGTTCTGGTG<br>CTGCTGACCAAAGCGGACAAACTGGCAAGCGGCGCACGTAAAGCGCAATTGAATAT<br>GGTGCGTGAAGCTGTACTGGCGTTTAACGGTGATGTGCAGGTTGAAACGTTTTCTTCG<br>TTGAAGAAACAAGGCGTGGACAAGCTGCGGCAGAAACTGGATACCTGGTTTAGCGAG<br>ATGCAGCCTGTAGAAGAAACGCAGGACGGCGAATAAAAGCTTGGCTGTTTT |
| **AP_GF33** | AGCGAATTCGAGCTCAGCAGGAGGAATTCAATGCGTGCTTTACCGATCTGTTTAGTAG<br>CACTCATGCTAAGCGGCTGTTCCATGTTAAGCAGATCCCCTGTCGAACCCGTTCAAA<br>GCACTGCACCCCAGCCGAAAGCGGAGCCTGCAAAACCGAAAGCGCCGCGCGCCA<br>CGCCGGTCCGAATTTATACCAATGCAGAAGAATTAGTCGGCAAACCGTTCCGCGATC<br>TCGGTGAAGTCAGTGGCGACTCTTGCCAGGCCTCTAATCAGGACTCTCCGCCGAGC<br>ATTCCAACCGCACGTAAGCGGATGCAAATCAACGCCTCTAAAATGAAAGCCAATGCT<br>GTATTACTGCATAGCTGCGAAGTCACCAGCGGTACGCCAGGCTGCTATCGTCAGGCT<br>GTATGTATCGGTTCTGCGCTTAACATTACGGCGAAATGAAAGCTTGGCTGTTTT |

| | |
|---|---|
| **AP_GJ1** | CGCCAATCGATATAAAGCAGGAGGAATTCAATGCAGTATTGGGGAAAAATCATTGGCG<br>TGGCCGTGGCCTTACTGATGGGCGGCGGCTTTTGGGGCGTAGTGTTAGGCCTGTTAA<br>TTGGCCATATGTTTGATAAAGCCCGTAGCCGTAAAATGGCGTGGTTCGCCAACCAGC<br>GTGAGCGTCAGGCGCTGTTTTTTGCCACCACTTTTGAAGTGATGGGGCATTTAACCAA<br>ATCCAAAGGTCGCGTCACGGAGGCTGATATTCATATCGCCAGCCAGTTGATGGACCG<br>AATGAATCTTCATGGCGCTTCCCGTACTGCGGCGCAAAATGCGTTCCGGGTGGGAAA<br>ATCAGACAATTACCCGCTGCGCGAAAAGATGCGCCAGTTTCGCAGTGTCTGCTTTGG<br>TCGTTTTGACTTAATTCGTATGTTTCTGGAGATCCAGATTCAGGCGGCGTTTGCTGATG<br>GTTCACTGCACCCGAATGAACGGGCGGTGCTGTATGTCATTGCAGAAGAATTAGGGA<br>TCTCCCGCGCTCAGTTTGACCAGTTTTTGCGCATGATGCAGGGCGGTGCACAGTTTG<br>GCGGCGGTTATCAGCAGCAAACTGGCGGTGGTAACTGGCAGCAAGCGCAGCGTGG<br>CCCAACGCTGGAAGATGCCTGTAATGTGCTGGGCGTGAAGCCGACGGATGATGCGA<br>CCACCATCAAACGTGCCTACCGTAAGCTGATGAGTGAACACCATCCCGATAAGCTGG<br>TGGCGAAAGGTTTGCCGCCTGAGATGATGGAGATGGCGAAGCAGAAAGCGCAGGAA<br>ATTCAGCAGGCATATGAGCTGATAAAGCAGCAGAAAGGGTTTAAATGACTAAGTAATAT<br>GGTG |
| **AP_GJMF** | CGCCAATCGATATAAAGCAGGAGGAATTCAATGGATGTCATTAAAAAGAAACATTGGTG<br>GCAAAGCGACGCGCTGAAATGGTCAGTGCTAGGTCTGCTCGGCCTGCTGGTGGGTT<br>ACCTTGTTGTTTTAATGTACGCACAAGGGGAATACCTGTTCGCCATTACCACGCTGATA<br>TTGAGTTCAGCGGGGCTGTATATGTTTGATAAAGCCCGTAGCCGTAAAATGGCGTGGT<br>TCGCCAACCAGCGTGAGCGTCAGGCGCTGTTTTTTGCCACCACTTTTGAAGTGATGG<br>GGCATTTAACCAAATCCAAAGGTCGCGTCACGGAGGCTGATATTCATATCGCCAGCC<br>AGTTGATGGACCGAATGAATCTTCATGGCGCTTCCCGTACTGCGGCGCAAAATGCGT<br>TCCGGGTGGGAAAATCAGACAATTACCCGCTGCGCGAAAAGATGCGCCAGTTTCGC<br>AGTGTCTGCTTTGGTCGTTTTGACTTAATTCGTATGTTTCTGGAGATCCAGATTCAGGCG<br>GCGTTTGCTGATGGTTCACTGCACCCGAATGAACGGGCGGTGCTGTATGTCATTGCA<br>GAAGAATTAGGGATCTCCCGCGCTCAGTTTGACCAGTTTTTGCGCATGATGCAGGGC<br>GGTGCACAGTTTGGCGGCGGTTATCAGCAGCAAACTGGCGGTGGTAACTGGCAGCA<br>AGCGCAGCGTGGCCCAACGCTGGAAGATGCCTGTAATGTGCTGGGCGTGAAGCCG<br>ACGGATGATGCGACCACCATCAAACGTGCCTACCGTAAGCTGATGAGTGAACACCAT<br>CCCGATAAGCTGGTGGCGAAAGGTTTGCCGCCTGAGATGATGGAGATGGCGAAGCA<br>GAAAGCGCAGGAAATTCAGCAGGCATATGAGCTGATAAAGCAGCAGAAAGGGTTTAA<br>ATGACTAAGTAATATGGTG |

**Supplementary References:**


1. Baba T, Ara T, Hasegawa M, Takai Y, Okumura Y, Baba M, et al. Construction of Escherichia coli K-12 in-frame, single-gene knockout mutants: the Keio collection. Mol Syst Biol. 2006;2:2006.0008. Epub 20060221. doi: 10.1038/msb4100050. PubMed PMID: 16738554; PubMed Central PMCID: PMCPMC1681482.
2. Karimova G, Pidoux J, Ullmann A, Ladant D. A bacterial two-hybrid system based on a reconstituted signal transduction pathway. Proc Natl Acad Sci U S A. 1998;95(10):5752-6. doi: 10.1073/pnas.95.10.5752. PubMed PMID: 9576956; PubMed Central PMCID: PMCPMC20451.
3. Brill JA, Quinlan-Walshe C, Gottesman S. Fine-structure mapping and identification of two regulators of capsule synthesis in Escherichia coli K-12. J Bacteriol. 1988;170(6):2599-611. doi: 10.1128/jb.170.6.2599-2611.1988. PubMed PMID: 2836365; PubMed Central PMCID: PMCPMC211177.
4. Majdalani N, Hernandez D, Gottesman S. Regulation and mode of action of the second small RNA activator of RpoS translation, RprA. Mol Microbiol. 2002;46(3):813-26. doi: 10.1046/j.1365-2958.2002.03203.x. PubMed PMID: 12410838.
5. Majdalani N, Heck M, Stout V, Gottesman S. Role of RcsF in signaling to the Rcs phosphorelay pathway in Escherichia coli. J Bacteriol. 2005;187(19):6770-8. doi: 10.1128/jb.187.19.6770-6778.2005. PubMed PMID: 16166540; PubMed Central PMCID: PMCPMC1251585.
6. Cabrera JE, Jin DJ. Growth phase and growth rate regulation of the rapA gene, encoding the RNA polymerase-associated protein RapA in Escherichia coli. J Bacteriol. 2001;183(20):6126-34. doi: 10.1128/jb.183.20.6126-6134.2001. PubMed PMID: 11567013; PubMed Central PMCID: PMCPMC99692.
7. Sharan SK, Thomason LC, Kuznetsov SG, Court DL. Recombineering: a homologous recombination-based method of genetic engineering. Nat Protoc. 2009;4(2):206-23. doi: 10.1038/nprot.2008.227. PubMed PMID: 19180090; PubMed Central PMCID: PMCPMC2790811.
8. Svenningsen SL, Costantino N, Court DL, Adhya S. On the role of Cro in lambda prophage induction. Proc Natl Acad Sci U S A. 2005;102(12):4465-9. Epub 20050223. doi: 10.1073/pnas.0409839102. PubMed PMID: 15728734; PubMed Central PMCID: PMCPMC555511.
9. Yahashiri A, Babor JT, Anwar AL, Bezy RP, Piette EW, Arends SJR, et al. DrpB (YedR) Is a Nonessential Cell Division Protein in Escherichia coli. J Bacteriol. 2020;202(23). Epub 20201104. doi: 10.1128/jb.00284-20. PubMed PMID: 32900831; PubMed Central PMCID: PMCPMC7648144.
10. Wall EA, Majdalani N, Gottesman S. IgaA negatively regulates the Rcs Phosphorelay via contact with the RcsD Phosphotransfer Protein. PLoS Genet. 2020;16(7):e1008610. Epub 20200727. doi: 10.1371/journal.pgen.1008610. PubMed PMID: 32716926; PubMed Central PMCID: PMCPMC7418988.
11. Thompson KM, Rhodius VA, Gottesman S. SigmaE regulates and is regulated by a small RNA in Escherichia coli. J Bacteriol. 2007;189(11):4243-56. Epub 20070406. doi: 10.1128/jb.00020-07. PubMed PMID: 17416652; PubMed Central PMCID: PMCPMC1913397.
12. Battesti A, Tsegaye YM, Packer DG, Majdalani N, Gottesman S. H-NS regulation of IraD and IraM antiadaptors for control of RpoS degradation. J Bacteriol. 2012;194(10):2470-8. Epub 20120309. doi: 10.1128/jb.00132-12. PubMed PMID: 22408168; PubMed Central PMCID: PMCPMC3347191.

13. Gibson DG, Young L, Chuang RY, Venter JC, Hutchison CA, 3rd, Smith HO. Enzymatic assembly of DNA molecules up to several hundred kilobases. Nat Methods. 2009;6(5):343-5. Epub 20090412. doi: 10.1038/nmeth.1318. PubMed PMID: 19363495.
14. Guzman LM, Belin D, Carson MJ, Beckwith J. Tight regulation, modulation, and high-level expression by vectors containing the arabinose PBAD promoter. J Bacteriol. 1995;177(14):4121-30. doi: 10.1128/jb.177.14.4121-4130.1995. PubMed PMID: 7608087; PubMed Central PMCID: PMCPMC177145.
15. Cherepanov PP, Wackernagel W. Gene disruption in Escherichia coli: TcR and KmR cassettes with the option of Flp-catalyzed excision of the antibiotic-resistance determinant. Gene. 1995;158(1):9-14. doi: 10.1016/0378-1119(95)00193-a. PubMed PMID: 7789817.
16. Karimova G, Ullmann A, Ladant D. Protein-protein interaction between Bacillus stearothermophilus tyrosyl-tRNA synthetase subdomains revealed by a bacterial two-hybrid system. J Mol Microbiol Biotechnol. 2001;3(1):73-82. PubMed PMID: 11200232.
17. Karimova G, Dautin N, Ladant D. Interaction network among Escherichia coli membrane proteins involved in cell division as revealed by bacterial two-hybrid analysis. J Bacteriol. 2005;187(7):2233-43. doi: 10.1128/jb.187.7.2233-2243.2005. PubMed PMID: 15774864; PubMed Central PMCID: PMCPMC1065216.
18. Clarke DJ, Holland IB, Jacq A. Point mutations in the transmembrane domain of DjlA, a membrane-linked DnaJ-like protein, abolish its function in promoting colanic acid production via the Rcs signal transduction pathway. Mol Microbiol. 1997;25(5):933-44. doi: 10.1111/j.1365-2958.1997.mmi528.x. PubMed PMID: 9364918.
19. Guillier M, Gottesman S. Remodelling of the Escherichia coli outer membrane by two small regulatory RNAs. Mol Microbiol. 2006;59(1):231-47. doi: 10.1111/j.1365-2958.2005.04929.x. PubMed PMID: 16359331.
20. Mandin P, Gottesman S. Integrating anaerobic/aerobic sensing and the general stress response through the ArcZ small RNA. Embo j. 2010;29(18):3094-107. Epub 20100803. doi: 10.1038/emboj.2010.179. PubMed PMID: 20683441; PubMed Central PMCID: PMCPMC2944060.
